# Whole brain mapping of spinal-projecting neurons in larval zebrafish

**DOI:** 10.64898/2026.05.20.726602

**Authors:** Martin Carbo-Tano, Kevin Fidelin, Tom Welch, Sujatha Narayan, Misha Ahrens, Réjean Dubuc, Claire Wyart

## Abstract

To elicit voluntary movements and integrative reflexes underlying behavior, the brain sends command signals to the spinal cord via specialized long-range descending neurons, known as spinal-projecting neurons (SPNs). The vast and widespread distribution of SPNs, combined with their complex long-distance connectivity, poses a significant challenge for mapping their anatomical organization and associating specific populations with distinct functions. Here we took advantage of the transparency and genetic accessibility of larval zebrafish to uncover the fundamental principles of SPN anatomical organization in a Teleost. Using an optical backfilling method relying on photoactivable GFP, we generate a whole-brain map of all neurons sending axons towards the spinal cord. This approach reveals far more SPNs than previously described through conventional strategies, offering an unparalleled opportunity to revisit distinct spinal-projecting nuclei distributed across hindbrain, midbrain, and diencephalic structures. Combining information on cell location, morphology, and projection patterns, we propose tentative homological designations for zebrafish of SPN nuclei based on established descriptions in mammals and other vertebrates.

## Introduction

Although often studied as separate entities, the brain and spinal cord function in synergy to govern numerous bodily functions. The brain contains multiple functionally-distinct neuronal populations known as spinal-projecting neurons (SPNs) that project through various regions of the spinal cord, terminating in specific segments along the rostrocaudal axis (Liang et al., 2016; Nyberg-Hansen, 1965; Wolters et al., 1982). These descending pathways orchestrate a wide array of motor, autonomic, and sensory functions (Kuypers, 2011, 1964; Winter et al., 2025). While some SPNs can carry motor commands ranging from basic locomotion to sophisticated fine movements involving muscles in the head or throughout the body (Kim et al., 2017; Leiras et al., 2022; Wang et al., 2017), other SPNs modulate the autonomic system regulating vital processes such as thermoregulation, breathing, cardiovascular function, and metabolism (Swanson and Sawchenko, 1980; Verstegen et al., 2017). Additionally, numerous SPNs modulate the sensory detection of tactile, thermal, proprioceptive, and painful stimuli (François et al., 2017; Liu et al., 2018).

The location and funicular trajectories of SPN nuclei have been well-documented via tract-tracing across many vertebrate taxa, including mammals, birds, reptiles, amphibians, lungfishes, elasmobranchs, and cyclostomi (Adli et al., 1999; Cabot et al., 1982; Cruce and Newman, 1981; Crutcher et al., 1978; Hermann et al., 2003; Lacroix-Ouellette and Dubuc, 2023; Ronan, 1989; Ronan and Northcutt, 1985; Sánchez-Camacho et al., 2001; Smeets and Timerick, 1981a; Ten Donkelaar, 1976; ten Donkelaar et al., 1980a, 1981a; Ten Donkelaar, 1982a). Comparative anatomical studies consistently demonstrate that the anatomical distribution of descending systems in early-derived vertebrates rivals that of mammals (Cruce and Newman, 1984; Ryczko et al., 2016). However, a significant gap remains in our understanding of the anatomy of these descending centers and their contribution to integrated motor and physiological responses.

The larval zebrafish (*Danio rerio*) has proven to be an ideal model for bridging this gap, offering a clear window into evolutionary conservation (Carbo-Tano et al., 2023) due to its unique experimental accessibility to its brain organization (Burgess and Burton, 2023; Pansera et al., 2025). Yet, in this genetic model organism the identity and characteristics of descending SPNs are not fully known. While horseradish peroxidase (HRP) tracing in various teleost species has shown a remarkable similarity to mammalian distributions (Bosch and Roberts, 1994, 2001; Hlavacek et al., 1984; New et al., 1998; Oka et al., 1986; Prasada Rao et al., 1987a), early anatomical studies in larval zebrafish predominantly labeled a very small (< 200) subset of neurons characterized by large cell bodies and large-caliber axons running in the medial longitudinal fasciculus (Kimmel, 1982; Kimmel et al., 1982). The distinctive morphology and stereotyped positions of these cells allowed for a systematic classification across individuals with names for distinct neurons identified according to its dorsoventral and anteroposterior coordinates (Kimmel, 1982; Kimmel et al., 1982; Lee et al., 1993; Lee and Eaton, 1991a). These early identifications of SPNs based on retrograde labeling suggested that the zebrafish descending system was possibly simpler in comparison to other vertebrates (Cruce and Newman, 1984). Retrograde labeling typically performed by injections in segment 14-15 highlighted primarily large reticulospinal neurons in the caudal midbrain and pontine region, with only sparse labeling in the medulla leading to a focus on these neurons for subsequent functional investigations (Collins et al., 2025; Gahtan and O’Malley, 2003; Huang et al., 2013; Lau et al., 2025; Orger et al., 2008).

Subsequent studies utilized photoconvertible proteins such as Kaede that change their emission spectrum from green to red upon UV light exposure (Ando et al., 2002). By illuminating axons in the spinal cord with UV light, the photoconverted Kaede diffuses retrogradely to cell bodies, allowing for ratiometric differentiation between spinal-projecting (red) versus non-spinal-projecting (green) neurons. By expressing Kaede in genetically-defined neuronal populations, such optical backfills of neurons projecting to the spinal cord have revealed within the population of neurons expressing *vsx2* (former *chx10*) transcription factor, that the medulla of larval zebrafish contains many more reticulospinal neurons than previously thought (Kimura et al., 2013, 2006; Pujala and Koyama, 2019). Yet, these studies were limited to discrete, genetically identifiable neuronal subsets, leaving homological relationships across vertebrates unaddressed.

To label virtually all supraspinal neurons, we relied here on the photo-activable GFP (PAGFP) expressed pan neuronally. To map all spinal-projecting neurons in larval zebrafish, PAGFP can be activated using two photon illumination in a confined section of the spinal cord (Bianco et al., 2012; Ruta et al., 2010). This approach reveals far more neurons than conventional retrograde tracing strategies have previously identified, offering an unparalleled opportunity to revisit distinct spinal-projecting nuclei distributed across the hindbrain, midbrain, and diencephalic structures. By combining information on cell location, morphology, and projection patterns, we propose tentative anatomical homologies between these nuclei in fish and mammals. These proposed homologies will undoubtedly be refined by future studies integrating molecular, genetic, and functional data.

## Methods

### Animal care and transgenic lines

Animal handling and procedures were validated by the Paris Brain Institute (ICM) and the French National Ethics Committee (Comité National de Réflexion Éthique sur l’Expérimentation Animale; APAFIS no. 2018071217081175 and #51743-2024102515029150) in agreement with European Union legislation. To avoid pigmentation, all experiments were performed on *Danio rerio* larvae of AB background with the *mitfa*^−/−^ mutation. Adult zebrafish were reared at a maximum density of eight animals per liter in a 14/10 h light–dark cycle environment at 28.5 °C. Larval zebrafish were typically raised in Petri dishes filled with system water under the same conditions in terms of temperature and lighting as for adults. Transgenic larvae expressing PAGFP under the alpha-tubulin promoter used for the photoconversion experiments were raised in complete darkness to minimize global photoactivation at the early stages. Experiments were performed at 20 °C on animals aged between 5 and 6 days post fertilization (dpf). We used the double transgenic line referred to as *Tg(Cau.Tuba1:PAGFP; elavl3:FusionRed)* for simplicity throughout, by crossing *Tg(Cau.Tuba1:c3paGFP)^a7437^* with *Tg(elavl3:stGtACR1-FusionRed)* and screening for double-positive embryos. For anatomical interrogations, we used the double transgenic line *Tg(oxt:GAL4-VP16:UAS:mCherry)* referred to as *Tg(oxt:GAL4;UAS:mCherry)* by crossing *Tg(oxt:GAL4-VP16)^zf3181Tg^* (Wee et al., 2019) with *Tg(UAS:Eco.NfsB-mCherry)^rw0144Tg^* (Agetsuma et al., 2010) and *Tg(GnRH2:EGFP)* (Xia et al., 2014).

### Design of the Tg(elavl3:stGtACR1-FusionRed) stable line

To generate the *Tg(elavl3:stGtACR1-FusionRed)* transgenic line, the stGtACR1-FusionRed coding sequence was subcloned from pAAV-CKIIa-stGtACR1-FusionRed (Addgene, plasmid #105679) into a Tol2 destination vector under the control of the pan-neuronal elavl3 promoter. Transgenic zebrafish were generated using the Tol2 transposon-based method (White et al., 2008a); the construct was co-injected with Tol2 transposase mRNA into one-cell stage zebrafish embryos in a *casper* background (Urasaki et al., 2008; White et al., 2008b). Injected fish (F0) were raised to adulthood and outcrossed to *casper* fish to identify germline-transmitting founders. Positive F1 offspring were identified by screening for FusionRed fluorescence, and stable transgenic lines were established from confirmed carriers.

### Optical backfills

Transgenic *Tg(Cau.Tuba1:PAGFP; elavl3:FusionRed) z*ebrafish larvae expressing cytosolic PAGFP and membrane tagged FusionRed proteins pan-neurally under the alpha-tubulin and elavl3 promoter respectively were raised in darkness. At 5 dpf, larval zebrafish were anesthetized in 0.02% MS-222 and mounted in glass-bottom dishes (MatTek, Ashland, MA, USA) dorsal side up in 1.5% low melting point agarose. The dish was filled with external bath physiological solution (134 mM NaCl, 2.9 mM KCl, 2.1 mM CaCl_2_-H_2_O, 1.2 mM MgCl_2_, 10 mM glucose and 10 mM HEPES, pH adjusted to 7.4 and osmolarity to 290 mOsm). The PAGFP protein in the rostral spinal cord (segments 4–9) was photoactivated in a two-photon microscope (2p-vivo, Intelligent Imaging Innovations, Inc., Denver, CO, USA, https://www.intelligent-imaging.com/) including a Mai Tai Deepsee laser (Spectra-Physics) tuned to 800 nm and a Zeiss Axio-Examiner Z1 with a Zeiss W Plan-APO 20X water-immersion objective (NA = 1). Line scans were controlled using Slidebook software (3i Intelligent Imaging Innovations, Inc., Denver, CO, USA). Before the photoactivation session, using minimum power the region of the spinal cord was recognized, and a region of interest was drawn in such a way that only half of the spinal cord was imaged. The photoactivation was done by scanning at 2 Hz with 4 µs dwell time using a laser power of 1 mW. The full depth of the spinal cord was scanned at a 1 µm interval and each plane was imaged for approximately 2 seconds. After the photoactivation session, larvae were unmounted and kept in the dark for 12 - 24h.

### Retrograde labeling experiments using biocytin

For the retrograde labeling experiments, 5-dpf *Tg(GnRH2:EGFP)* larvae were anesthetized with 0.02% MS-222 (catalog no. A5040, Sigma-Aldrich) and laterally embedded in 1.5% low melting point agarose (Sigma, A4018) prepared with external bath solution (as described above). A small portion of the agarose was removed on top of the tail around myomere 9, and all the liquid was then extracted from the dish. A few crystals of biocytin Alexa Fluor 594 (catalog no. A12922, Thermo Fisher Scientific) were diluted in 5 µl distilled water and then allowed to recrystallize on the tip of insect pins (stainless steel Minutiens, 0.2-mm diameter, Austerlitz). Under a stereo dissecting microscope, a unilateral single lesion in the spinal cord between myomere 8 and myomere 10 was done using the dye-soaked insect pins. After the procedure, the dish was filled with external solution, the larvae were unmounted and allowed to recover for 2 h in an oxygenated external solution. Larvae were allowed to recover overnight in an external bath solution. The next day, the larvae were anesthetized again with 0.02% MS-222 and mounted dorsally in 1.5% low-melting-point agarose. The retrogradely labeled neurons were imaged in a confocal microscope as described above.

### Whole-brain imaging

At 6 dpf, larval zebrafish were anesthetized in 0.02% MS-222 (catalog no. A5040, Sigma-Aldrich) and mounted in glass-bottom dishes dorsal side up in 1.5% low melting point agarose. The brain and rostral spinal cord were imaged in an upright confocal microscope (Olympus FV1200 confocal system, Shinjuku City, Tokyo, Japan) with a 20X water dipping objective NA = 0.5 (UMPLFLN 20XW) for capturing the general brain features and a 40X water dipping objective NA = 0.80 (LUMPLFLN 40XW) for further detailed imaging. Using the 20X objective, the whole brain of larval zebrafish was captured in two consecutive tiles with dimension 500 x 510 µm (10% overlap). Voxel size was set to ∼ 0.9 x 0.9 x 2 µm. Detailed imaging with the 40X objective was done with a voxel size of ∼ 0.2 x ∼0.2 x 2 µm. Due to the somewhat diffuse PAGFP signal when photoactivated, natural tissue autofluorescence when imaging with 488nm light, and because all imaging was performed from the dorsal aspect, the image quality is degraded in the ventral regions of the brain.

### Reference brain creation and 3D registration

To generate the reference brain space, the consecutive tiles of each brain were stitched using Fiji (i.e., ImageJ) software (https://fiji.sc/) applying the stitching plugin (Preibisch et al., 2009). Our template brain was a 6-dpf *Tg(elavl3:stGtACR1-FusionRed);mitfa^−/−^* larvae. Registration of image volumes was performed using ANTs (http://stnava.github.io/ANTs/) through the SlicerANTs extension (Revision: 4e622b5) of 3DSlicer software (Fedorov et al., 2012), www.slicer.org, version 5.8.1). The soma targeting of the opsin GtACR1 was not effective in the line *Tg(elavl3:stGtACR1-FusionRed)*, which enabled us to have a membrane signal providing a robust scaffold of the brain architecture. The FusionRed signal of 6 larvae was registered using affine and SyN registration steps. We created the mean-stack reference brain by calculating the mean across all registered fish and normalizing to the maximum-intensity. Registration steps followed and ANTs parameters used are described in (Bhandiwad et al., 2024). All registrations were manually assessed for global and local alignment accuracy. In order to align our results with the well-established effort of the mapZbrain (Kunst et al., 2019) and Zbrain (Randlett et al., 2015) atlases, we registered the *Tg(elavl3:lynTagRFP)* signal from the *mapZbrain* Atlas and Anti-Zrf2 signal from the *Zbrain* atlas was registered to our *Tg(elavl3:FusionRed)* space.

### Cell detection

Single-cell detection and segmentation were performed manually due to the complex nature of the backfill signal. Automatic segmentation methods failed to accurately identify individual neurons because the backfilled signal included not only somata but also dendrites and axons, resulting in a poor signal-to-noise ratio. Individual neurons were identified and manually segmented based on soma morphology and labeling intensity using *napari* software (Sofroniew et al., 2026) (https://github.com/napari/napari). Each neuron was annotated using the point layer tool in *napari*, with points placed at the center of each soma. Neurons were traced across consecutive optical sections to ensure accurate identification and avoid double-counting the same neuron. The spatial *xyz* coordinates of each identified neuron were recorded for subsequent anatomical registration and analysis.

### Data analysis and visualization

Analysis was done either in R (v.4.5.3) using the RStudio IDE (v.2025.9.0.387) (Posit team, 2025) or Python v.3.7. Figures were assembled in Inkscape (v.1.4.3, https://inkscape.org/).

## RESULTS

### Brainwide distribution of spinal descending neurons

To investigate the distribution of neurons sending descending axons to the spinal cord in larval zebrafish, we employed optical backfilling in 5-day post fertilisation (dpf) double transgenic larvae *Tg(Cau.Tuba1:PAGFP; elavl3:FusionRed)*. These transgenic larvae pan-neuronally express cytosolic photoactivatable GFP (PAGFP) and membrane-targeted FusionRed (**Figure 1a**). We performed unilateral photoactivation at the rostral spinal cord using a two-photon scanning system at 800 nm on a region extending caudally from the obex and spanning approximately 10 spinal segments (**Figure 1a, Methods**). At 6-dpf, larvae were mounted in agarose and their brains were imaged using confocal microscopy (**Figure 1a**). To enable direct comparison of results across individual animals and to integrate our findings with existing publicly available zebrafish brain atlases, we generated an averaged brain template from six whole-brain images based on the FusionRed signal (**Figure 1b**). We then bridged our brain template with the well-established zebrafish atlases registering our brain template with the Zbrain Anti-Zrf2 signal (Randlett et al., 2015) and the mapZbrain *Tg*(*elavl3:lyn-TagRFP)* signal (Kunst et al., 2019). Following the registration to the template brain, PAGFP-positive cell bodies were manually segmented from the 3D image stacks. The resulting whole-brain distribution map revealed a substantial population of spinal-projecting neurons in larval zebrafish (**Figure 1c**, **Figure 2, Supp.video 1**), displaying a clear ipsilateral prominence with an average of 654.5 ± 53 (mean ± SD) spinal-projecting neurons per larva (estimated 1,309 ± 106 neurons for the complete spinal-projecting population (mean ± SD)). To regionalize the locations of the segmented neurons, we utilized the anatomical masks of the Zbrain atlas (Randlett et al., 2015), which provides better anatomical annotation in the hindbrain region where the rhombomeric boundaries are chosen (see below). Consistent with the distribution of spinal-projecting neurons in other vertebrates, we found that the hindbrain contains the largest population of spinal descending neurons (**Figure 1d**), with the medulla oblongata harboring the vast majority of these cells (**Figure 1c**). Some anatomical regions displayed a clear unilateral distribution pattern, while other neuronal populations were primarily bilateral in nature (**Figure 2c**).

**Figure 1.**
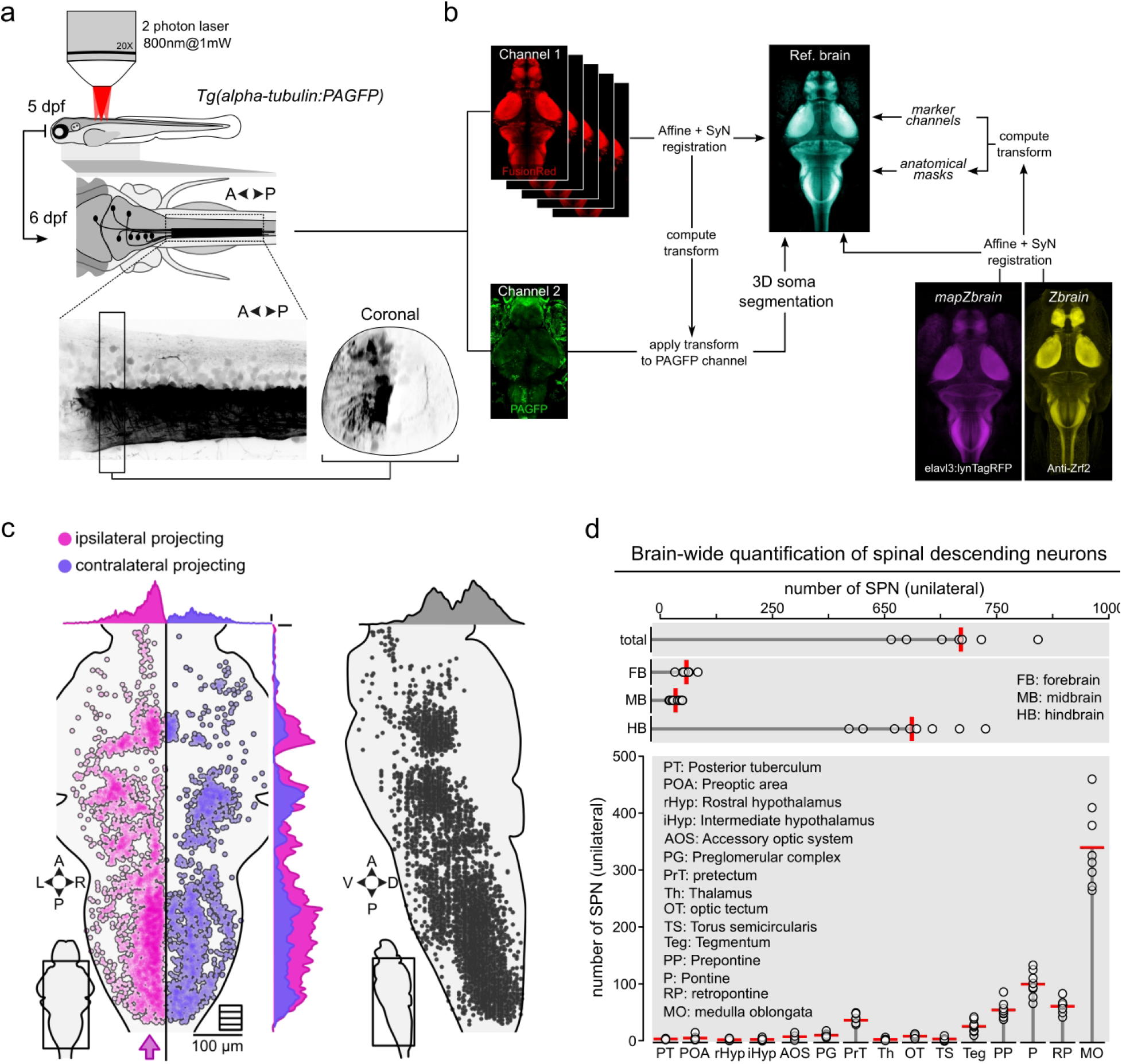
Whole-brain atlas of spinal-projecting neurons in larval zebrafish. **a.** Schematic illustrating the principle of optical backfilling using photoactivatable GFP to label spinal projection neurons. **b.** Image processing workflow for anatomical registration and alignment to a standard brain reference. **c.** Three-dimensional distribution of segmented soma locations of spinal projection neurons in 6-day post-fertilization (dpf) larval zebrafish. The arrow indicates the site of optical backfilling in the spinal cord. Neurons are color-coded according to projection laterality: ipsilateral (magenta) and contralateral (purple). **d.** Quantitative analysis of spinal-projecting neuron quantification across brain regions. The red line indicates the mean value and each circle represents a larva (n = 8 larvae).

**Figure 2.**
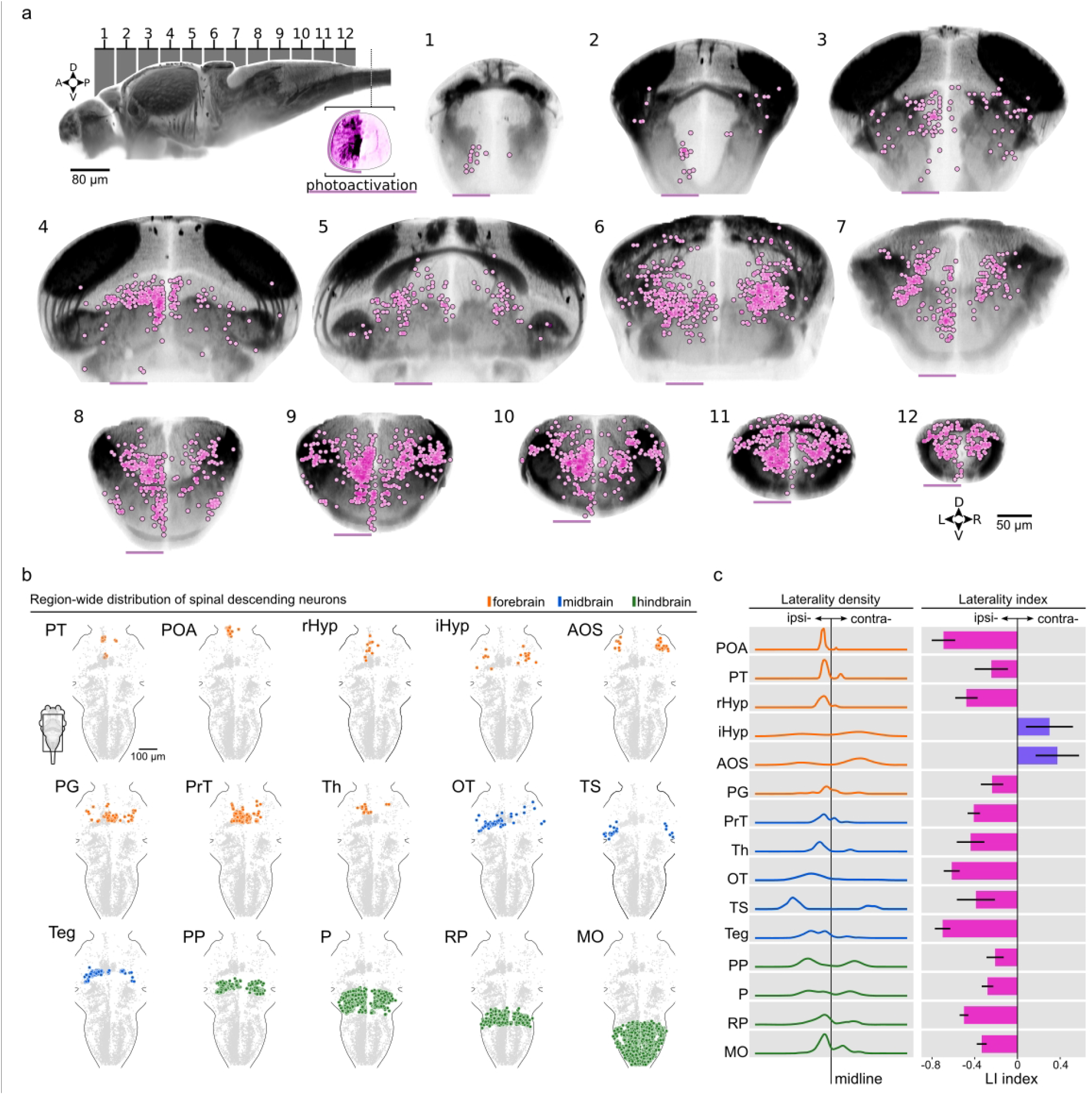
Regional distribution and laterality of spinal-projecting neurons in larval zebrafish. **a.** Coronal sections showing the spatial distribution of descending spinal projection neurons throughout the brain. The top left panel displays a lateral view of the reference brain atlas with lines indicating the anatomical position and thickness of each coronal section plane. Magenta lines below each coronal section denote the spinal cord location where optical backfilling was performed. **b.** Spatial organization of spinal projection neurons across anatomical regions defined by the Zbrain atlas. **c.** Density of projection laterality (Mean + SD, n = 8 larvae) (left panel) and laterality index (right panel) for across the anatomical regions studied. *LI= (N_contra - N_ipsi) / (N_ipsi + N_contra). LI* = Laterality Index, *N_ipsi* = Number of spinal projection neurons on the ipsilateral side and *N_contra* = Number of spinal projection neurons on the contralateral side. Anatomical abbreviations as in Figure 1d.

The identification and characterization of spinal-projecting neurons (SPNs) in zebrafish relied on multiple complementary criteria. The primary consideration was the spatial organization of labeled neurons, including both their absolute coordinates in the brain atlas and their relative positioning with respect to known nuclei and fiber tracts (medial longitudinal fasciculus, MLF; lateral longitudinal fasciculus, LLF; posterior cerebellar tract, PC; and vestibulo-spinal tract, TVS). Cellular morphology provided additional distinguishing features, particularly the characteristics of labeled cell bodies, including soma size, and density. When detectable, the trajectory patterns of axons emerging from labeled cell bodies served as another crucial identification criterion. We documented the distribution patterns, noting both symmetrical (reflecting ipsilateral and contralateral projections) and asymmetrical (reflecting either ipsilateral or contralateral projections) labeling. The relative intensity of labeling within cell groups provided supplementary discriminating information, as consistent differences in signal strength emerged between distinct neuronal populations. Unlike conventional retrograde labeling methods employing HRP or conjugated dextrans, optical backfilling reliably labels somatic compartments but does not yield sufficient signal to resolve dendritic or axonal morphology. We attribute this limitation to two factors: the simultaneous labeling of large numbers of neurons, which likely increases background fluorescence, and the restricted diffusion of photoactivated PAGFP into dendritic compartments. Therefore, all our subsequent analyses focus primarily on somatic information. Our analysis extended beyond the anatomical characteristics of nuclei to encompass broader organizational principles, as we anchored our observations with expression patterns from reference transgenic lines and hybridization chain reaction (HCR) staining data available in the Zbrain and mapZbrain atlases (https://zebrafishexplorer.zib.de/; https://mapzebrain.org/). In the following sections, we provide a detailed description of identifiable cellular groups of SPNs organized in rostrocaudal order.

### Telencephalic projections to spinal cord

Using classical tracing techniques, mammals appeared as the main taxon of vertebrate species with obvious telencephalic projections to the spinal cord (Cruce et al., 1999) along with salamanders and sharks where few telencephalic spinal projection neurons (Kokoros and Northcutt, 1977). In some of our specimens (n = 4 out of 8), few telencephalic neurons in the olfactory bulb sent axons to the spinal cord (**Figure 3a**). Using the ZExplorer atlas (Du et al., 2025), which contains thousands of single neuron tracings in the larval zebrafish brain, we searched for telencephalic neurons projecting to the spinal cord. We confirm that a few spinal-projecting neurons in the olfactory bulbs send both ipsilateral and contralateral descending axons, while in the subpallium spinal projection neurons send only ipsilateral descending axons (**Figure 3a**). The neurons we found in the olfactory bulbs matched those present in the ZExplorer atlas (**Figure 3a**). In contrast, we could not detect subpallial neurons in our dataset, likely because these neurons are very sparse or because the photoactivation level was insufficient to distinguish their somas from the background signal.

**Figure 3.**
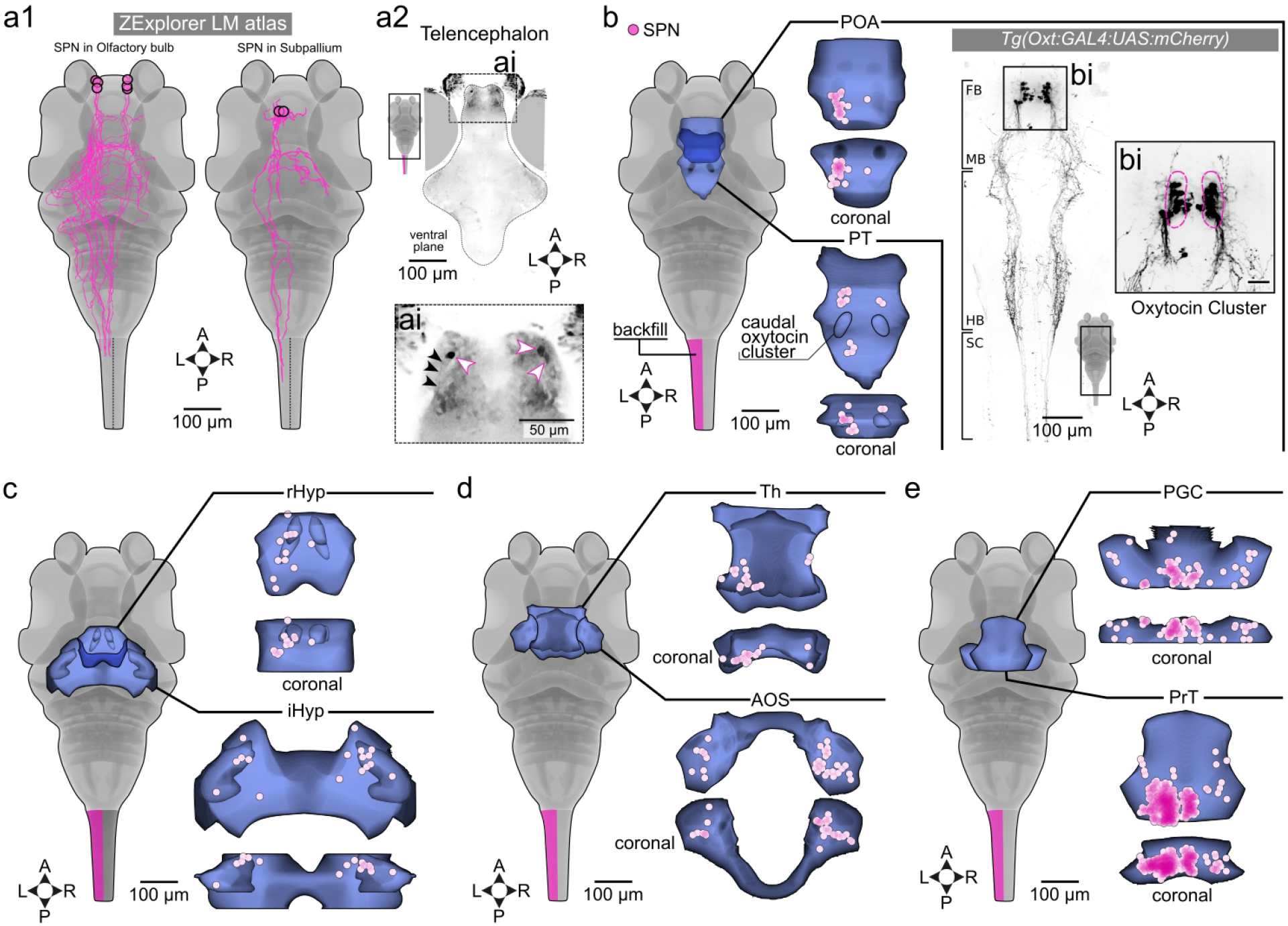
Telencephalic and diencephalic spinal-projecting neurons. **a1.** spinal-projecting neurons revealed using the ZExplorer larval zebrafish atlas. The web interface (https://zebrafish.cn/LM-Atlas/) was queried to identify neurons with somata in the telencephalon and axonal projections to the spinal cord. **a2.** Spinal projection neurons identified by optical backfilling in the olfactory bulb. White arrows indicate somata of ipsilateral and contralateral projecting neurons; black arrows trace the trajectory of a representative descending axon. **b.** Distribution of spinal projection neurons in the preoptic area (POA) and posterior tuberculum (PT). Neurons are displayed within their respective anatomical boundaries in horizontal (top) and coronal (bottom) orientations. The right panel shows localization and expression pattern of oxytocin-expressing neurons of the POA in the *Tg(oxt:GAL4;UAS:mCherry)* transgenic line. **bi.** Scale bar, 10 µm. **c.** Spinal projection neurons in the hypothalamus, subdivided into rostral (rHyp) and intermediate (iHyp) regions. **d.** Spinal projection neurons in the thalamus (Th) and accessory optic system (AOS). **e.** Spinal projection neurons in the preganglionic complex (PGC) and pretectum (PrT).

### Diencephalic projections

The preoptic area (POA) contains only ipsilateral projecting neurons (**Figure 3b**, **Figure 2c**). These neurons correspond to the rostral oxytocinergic cluster as shown by the expression pattern of the *Tg(oxt:GAL4;UAS:mCherry)* transgenic line (**Figure 3b**). In agreement with the results of our optical backfills, the *oxt+* neurons send exclusively ipsilateral descending axons to the spinal cord (**Figure 3b**). Caudally, the posterior tuberculum (PT) has also mostly ipsilateral projection neurons (**Figure 2c**) segregated into two clusters, one anterior and the other posterior to the caudal oxytocin cluster (**Figure 3b**). The location of these two spinal-projecting clusters corresponds to the diencephalic dopaminergic (DA) groups referred to as DC2 and DC4 in zebrafish (Tay et al., 2011), and is in line with the results found in lampreys and other vertebrates (Ryczko et al., 2016). In the hypothalamus, spinal-projecting neurons were localized in the rostral (rHyp) and intermediate subdivisions (iHyp). While spinal-projecting neurons in the rHyp are exclusively ipsilateral, those in the iHyp show a slight contralateral bias (**Figure 3c**, **Figure 2c**). The thalamus (Th) hosts mainly ipsilateral spinal-projecting neurons located in the caudal pole (**Figure 3d**, **Figure 2c**). We found many spinal-projecting neurons in the accessory optic system (AOS). The AOS sends bilateral projections to the spinal cord with a strong contralateral prominence (**Figure 3d**, **Figure 2c**). The preganglionic complex (PGC) contains the soma of many spinal-projecting neurons with ipsi- and contralateral projections to the spinal cord, with the highest density near the midline adjacent to the caudal pretectum (**Figure 3e**, **Figure 2c**). At this level, the pretectum (PrT) hosts the soma of many neurons projecting to the spinal cord that are densely packed along the midline with two easily recognizable nuclei, a large one with strong ipsilateral labeling and a smaller one located more medially with contralateral projections (**Figure 3e**, **Figure 2c**).

The PGC and the PrT are two closely located regions with boundaries that are difficult to define. The region comprising the caudal PGC, PrT, and rostral tegmentum (Teg) is also known as the interstitial tegmentum or central tegmental field (Ten Donkelaar, 2020; Warner, 1947), an ill-defined area with nuclei that project to large regions of the central nervous system (Walter and Shaikh, 2014). There is still confusion about whether these areas belong to the forebrain or the midbrain (Watson et al., 2019a). In most vertebrate species investigated, conspicuous spinal-projecting neurons have been described in this area (Auclair et al., 1993; Castiglioni et al., 1978; Kuypers and Maisky, 1975; Nudo and Masterton, 1988a; ten Donkelaar et al., 1980b; Ten Donkelaar, 1982b). In zebrafish, the spinal-projecting neurons in this region have been classically named the nucleus of the medial longitudinal fasciculus (nMLF) (Kimmel, 1982; Kimmel et al., 1982) and have been regarded as located in the midbrain (Berg et al., 2023). Nevertheless, recent reports have relocated the nMLF, using molecular approaches (Wullimann et al., 2023), to prosomere 1 of the diencephalon, aligning its location with the corresponding locus in mammals (Watson et al., 2019a). Since the anatomical masks used in this study still rely on the uncorrected region boundaries, we opted to assign the spinal-projecting neurons corresponding to the rostral medial tegmentum to the caudal PrT.

The pretectal interstitial nuclei are composed of the centrally projecting Edinger-Westphal nucleus (EW), the nucleus of Darkschewitsch (ND), and the interstitial nucleus of Cajal (INC) laterally (Ten Donkelaar, 2020) (**Figure 4a**). All three nuclei send projections to the spinal cord, unilaterally for the ND and INC, and with a bilateral tendency for the EW (Liang et al., 2011a; Nudo and Masterton, 1988b). In mice, the EW is a facultatively peptidergic nucleus that expresses numerous peptides, including Cocaine- and amphetamine-regulated transcript (CART) and Urocortin (Ucn) (Bluett et al., 2025; Priest et al., 2023). Anatomically, the centrally projecting EW has extensive projections throughout the brain and spinal cord (Topilko et al., 2022). In an attempt to find zebrafish transgenic lines useful for targeting the nMLF, we identified the *Tg(GnRH2:EGFP)* (Xia et al., 2014) transgenic line as a possible candidate (**Figure 4b**). At 6 dpf, the *Tg(GnRH2:EGFP)* transgenic line labels only a single cluster of neurons in the interstitial tegmental region with widespread projections throughout the brain and spinal cord (Xia et al., 2014; Yamamoto et al., 1995a). However, labeling neurons in the nMLF with spinal dye injections revealed that the GnRH2^+^ cluster lies medial to the nMLF (**Figure 4c1**) (Yamamoto et al., 1998). We previously proposed that the nMLF of zebrafish larvae might correspond to the INC (Carbo-Tano et al., 2023). Given that the *Tg(GnRH2:GFP)^+^* cluster is located medially to the INC and has an extensive projection pattern, we speculated that the *Tg(GnRH2:GFP)+* cluster corresponds to the EW nucleus. We tested this hypothesis by examining the expression pattern of two peptides associated with the EW nucleus in mammals, using the whole-brain distribution of mRNA for cocaine- and amphetamine-regulated transcript 2 (*cart2*) and urotensin 1 (*uts1*) from the mapZbrain atlas (homologs to CART and UCN in mammals). We found that the signals for *cart2* and *uts1* match the GnRH2^+^ cluster, supporting our hypothesis that the GnRH2^+^ cluster corresponds to the peptidergic EW nucleus (**Figure 4C2**). Using the EW as an anchor point, we now propose that the interstitial pretectal nuclei are composed of the EW medially, the ND adjacent laterally, and the INC lateral to the MLF (**Figure 4d**). Since the ND is included as part of the ventral paraqueductal gray (PAG) (Ingram and Ranson, 1935; Zhao et al., 2024), the spinal-projecting neurons of the ND in the most dorsal planes could be part of the ventral PAG, which is known to have ipsilateral projections to the spinal cord (Liang et al., 2011a).

**Figure 4.**
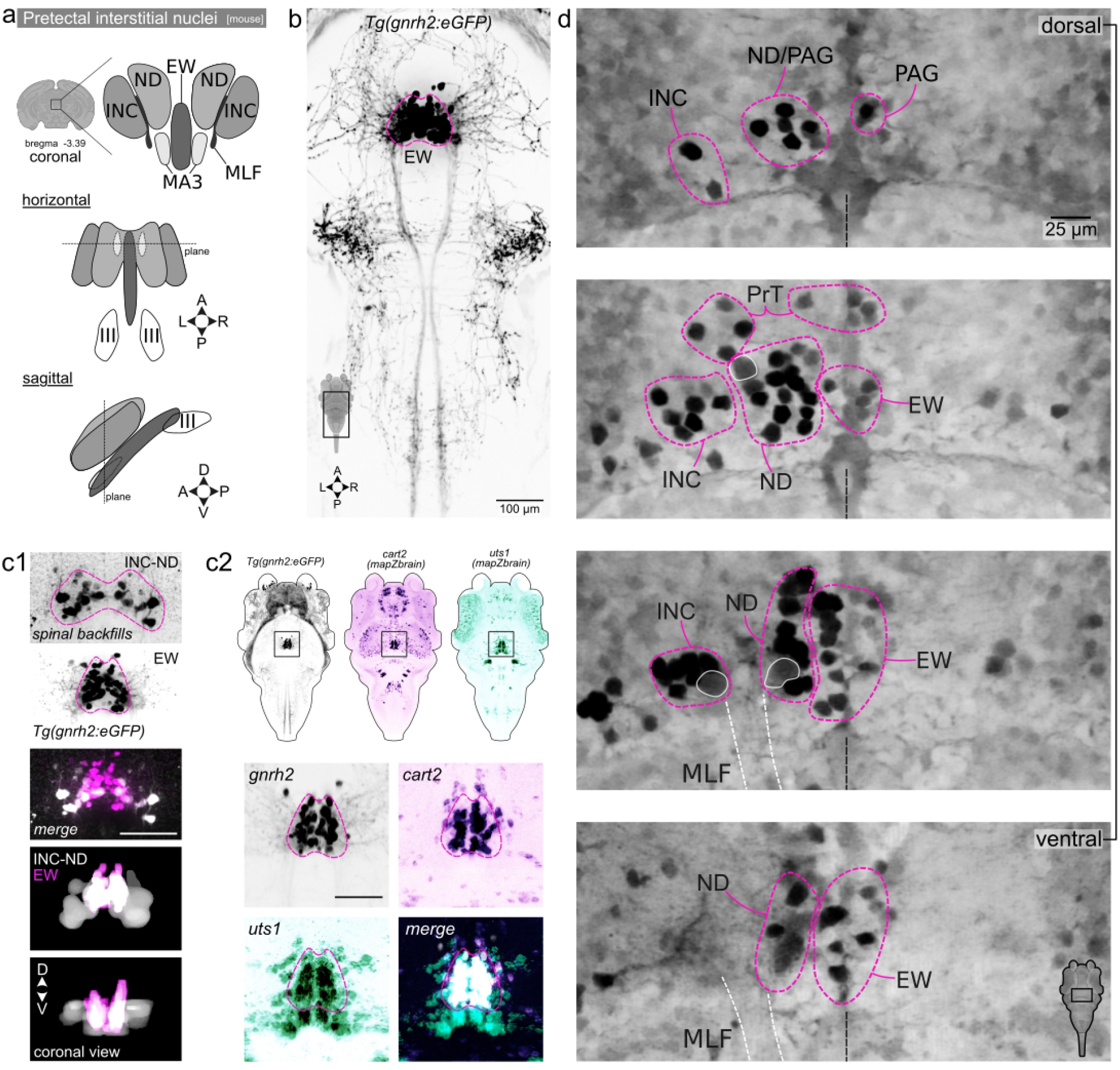
Identification of pretectal interstitial nuclei homologs in larval zebrafish. **a.** Schematic representation of the mammalian pretectal interstitial nuclei organization and composition. **b.** Spatial distribution of GnRH2-expressing neurons in the *Tg(gnrh2:eGFP)* transgenic line. **c1.** Pretectal Edinger-Westphal (EW) nucleus homolog gnrh2^+^ neurons are not labeled by classical retrograde spinal dye backfills. Lower panels show horizontal (top) and coronal (bottom) views of anatomical masks for the EW and the classically designated nucleus of the mediolateral fasciculus (nMLF) (putative Interstitial Nucleus of Cajal - nucleus of Darkschewitsch (INC-ND) homolog). The EW nucleus is positioned medially to the INC-ND nuclei labeled by spinal backfills. Scale bar 40 µm. **c2.** Anatomical comparison of pretectal GFP^+^ neurons in the *Tg(gnrh2:GFP)* line with the expression patterns of *cart2* and *uts1* mRNA from the mapZbrain atlas. **d.** Confocal image series through the pretectal interstitial region from ventral (bottom) to dorsal (top). Proposed homology assignments for zebrafish pretectal interstitial nuclei relative to mammalian counterparts are indicated. MLF, mediolateral fasciculus.

### Mesencephalic projections

In the mesencephalon, retrogradely-labeled neurons were observed in the optic tectum (OT) with an ipsilateral prominence (**Figure 5a**, **Figure 2c**). The contralaterally projecting neurons are few and sparse and not spatially organized into a cluster. The highest density of neurons is located ipsilateral to the backfill in the ventrocaudal periventricular layer in close proximity to the tegmentum (Teg). Retrogradely labeled neurons were also found in the torus semicircularis (TS) on both sides of the brain, but with a slight ipsilateral bias (**Figure 5b**).

**Figure 5.**
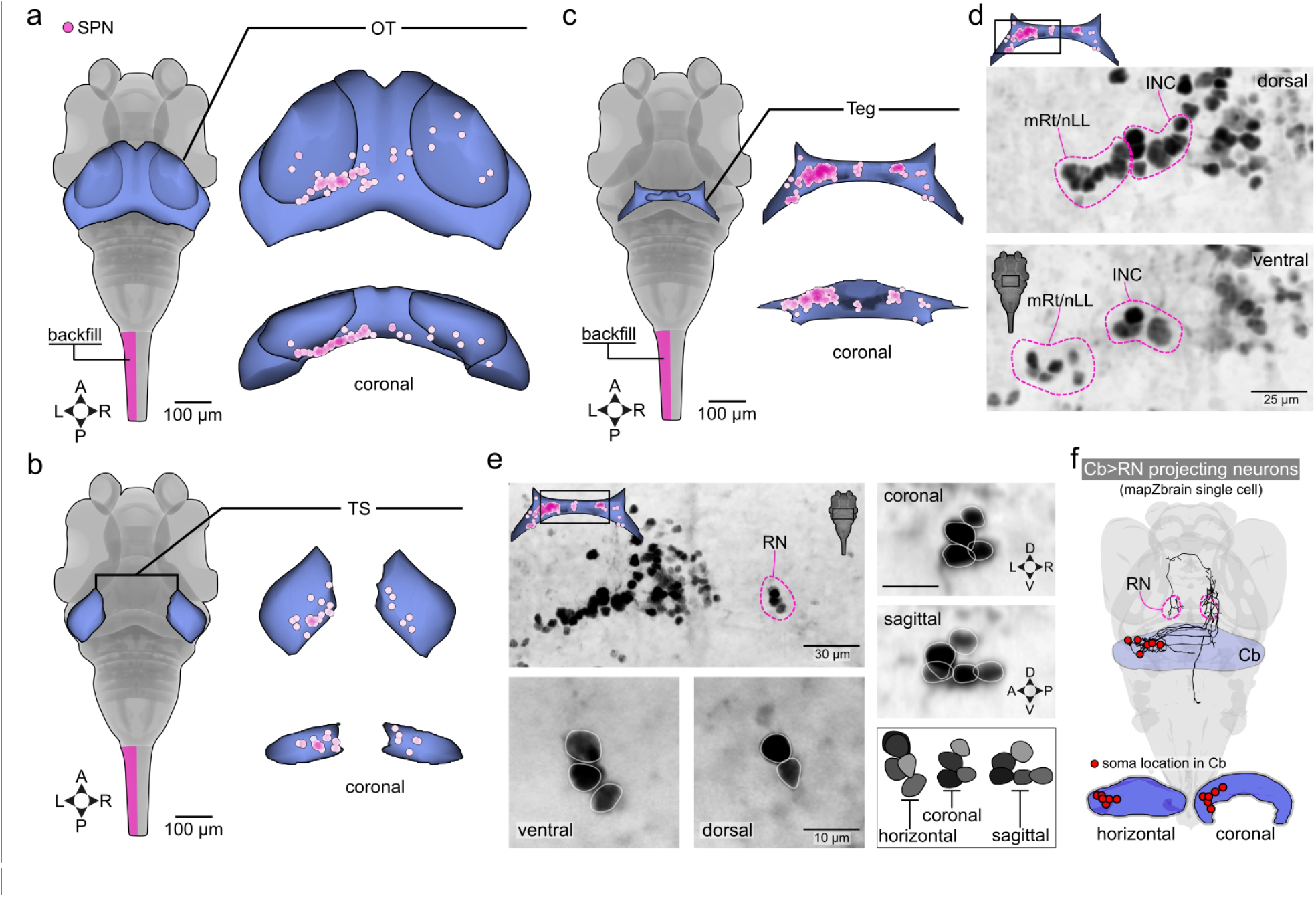
Mesencephalic spinal-projecting neurons. **a.** Distribution of spinal projection neurons in the optic tectum (OT). **b.** Distribution of spinal projection neurons in the torus semicircularis (TS). **c.** Distribution of spinal projection neurons in the tegmentum (Teg). d. Confocal image series through the mediolateral tegmentum (boxed region in inset) from ventral (bottom) to dorsal (top). Proposed homology assignments for zebrafish midbrain reticular formation (mRF) and nucleus of the lateral lemniscus (nLL) are indicated. **e.** Confocal images of the mid-tegmentum (boxed region in inset) showing the contralaterally projecting red nucleus (RN). Lower panels display ventral and dorsal aspects of the RN with constituent neurons delineated by white outlines. Right panels show coronal (top), sagittal (middle), and schematic (bottom) representations of RN neuronal organization. **f.** Single-cell projection patterns of cerebello-rubral neurons identified using the mapZbrain single-cell atlas (https://mapzebrain.org). The RN region was used as a seed to query neurons with axonal projections to this target area.

The Teg contains spinal-projecting neurons on both sides of the brain. At the midline, we found ipsilateral labeling of a group of neurons that could correspond to the medial accessory oculomotor nucleus (MA3) of mammals (Liang et al., 2011b). Lateral to the INC, we consistently found a group of well-labeled neurons sending ipsilateral axons to the spinal cord (**Figure 5c-d**). Based on their location and projection pattern, these neurons could belong to the midbrain reticular nucleus (mRt) or nucleus of the lateral lemniscus (nLL) as described in teleost (New et al., 1998; Yáñez et al., 2024) (**Figure 5c-d**). Sending exclusively contralateral projections, we found the red nucleus (RN) in the mediolateral Teg (**Figure 5e**). Consistent labeling of the RN was difficult to achieve, with RN neurons showing variable labeling intensity, possibly because the RN in 6 dpf larval zebrafish does not project far caudally beyond the obex region. We found that the RN consists of 5 neurons arranged in a C shape in the sagittal plane (**Figure 5e**). Unfortunately, due to the consistently low level of photoactivation, we were not able to trace the axons of the RN neurons to confirm their projection characteristics. To further confirm our identification of this group of neurons as the RN, we investigated whether the cerebellum (Cb) sends contralateral efferents to the area we identified as the RN (Finger, 1978). To verify Cb to RN connections, we used the mapZbrain single cell atlas to search for projections to the region we identified as the RN. Confirming our hypothesis, we found neurons with somas in the Cb sending axons that terminate or pass through the RN (**Figure 5f**).

### Rhombencephalic projections

From this point caudally, the number of labeled neurons increases considerably (**Figure 1d**). For the brainstem description, we followed the nomenclature and subdivisions proposed by Watson et al. 2019 (Watson et al., 2019a). Although originally developed for mammals, we found this nomenclature and the brainstem boundaries based on developmental gene expression to be well-suited for comparative studies across vertebrates (Bhandiwad et al., 2022; Carbo-Tano et al., 2023). The brainstem starts with the prepontine (PP) region, composed of the isthmus and rhombomere 1 (r1), with the trochlear nucleus (4N) lying at the rostral border of the isthmus and the locus coeruleus (LC) at the caudal pole of r1 (Watson et al., 2019a). In the PP region, we found spinal-projecting neurons on both sides with nearly equal laterality and a slight ipsilateral bias (**Figure 6a**, **Figure 2c**). At the ventral planes, we found large neurons with ipsilateral projecting axons that are part of the pontine oralis group of mammals (Liang et al., 2011a). At the medial level, the LC is strongly labeled bilaterally, lying at the caudal border of r1 (**Figure 6b**). Rostral to the LC, we found a group of spinal-projecting neurons with bilateral projections to the spinal cord and an ipsilateral prominence that resembles the parabrachial group of mammals (PB) (Liang et al., 2011a) (**Figure 6b**). In mice, at the level of the trochlear nucleus in the isthmic region, the cuneiform nucleus (CuF) shows ipsilateral projections to the spinal cord (Liang et al., 2011a). At the same isthmic level, the midbrain reticular formation (mRt) shows bilateral projections to the spinal cord (Liang et al., 2011a). In larval zebrafish, we found a similar topographical organization for the spinal-projecting neurons located rostrolaterally to the PB neurons, where both ipsilateral and contralateral projecting groups can be identified (**Figure 6b**). The ipsilateral projecting group could correspond to the cuneiform nucleus (CuF) or the caudal pole of the mRt, while the contralateral projecting group likely corresponds to mRt (**Figure 6b**). Rostral to the LC, labeled somata were present with an ipsilateral prominence, likely belonging to the pedunculotegmental nucleus (PTg) or/and the laterodorsal tegmental nucleus (LDT) (**Figure 6b**) in the location where we previously functionally identified the mesencephalic locomotor region (MLR, (Carbo-Tano et al., 2023)). Along the midline, we found faint PAGFP+ neurons located at the caudal pole of the superior raphe nucleus (SR) (**Figure 6b, d**). These raphe spinal-projecting neurons had an ipsilateral prominence but were difficult to label consistently. By performing bilateral optical backfills, the labeling intensity was enhanced, revealing the sublocation of the raphe spinal-projecting neurons in the caudal pole (**Figure 6d**).

**Figure 6.**
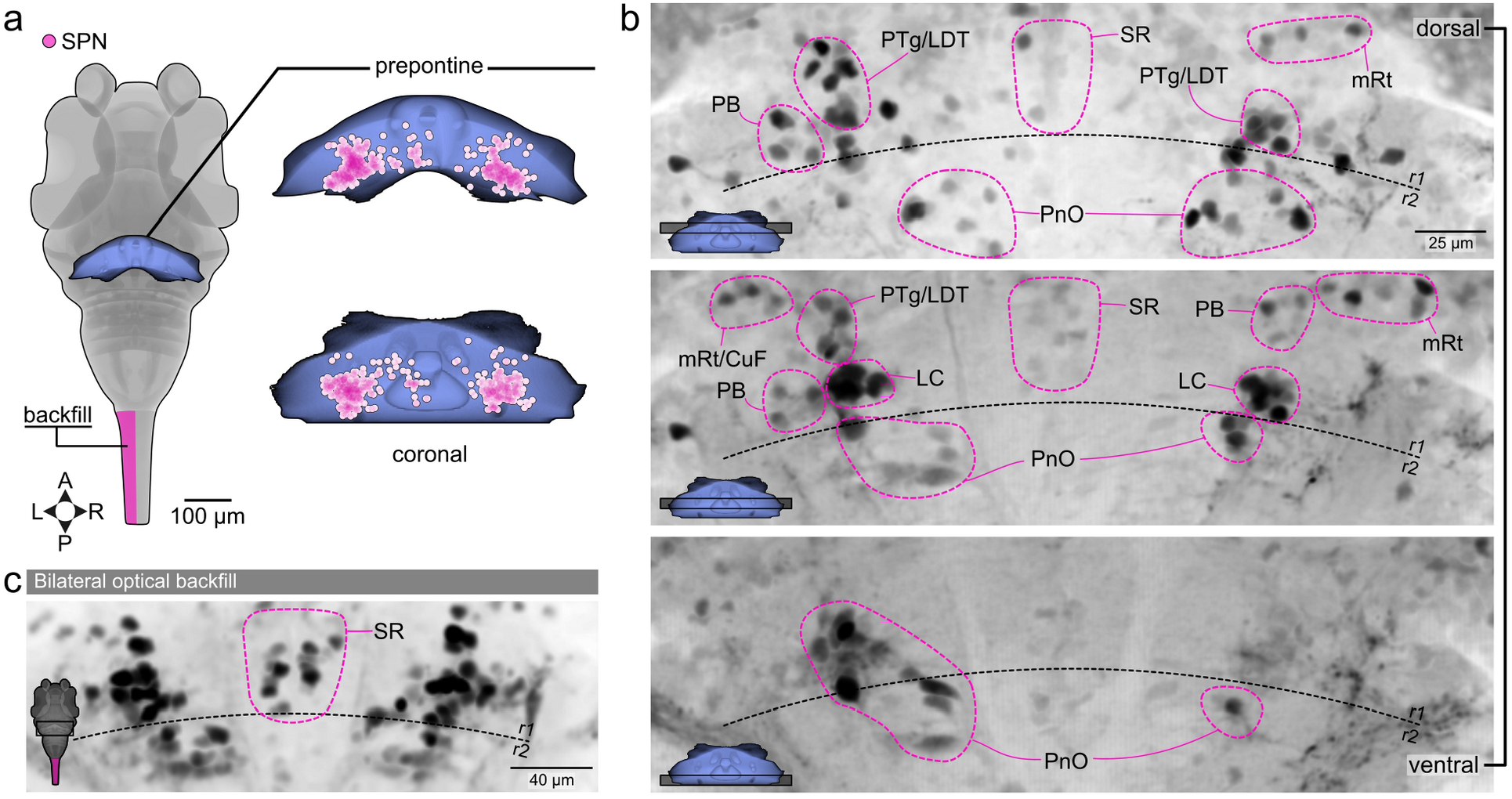
Rhombencephalic spinal-projecting neurons: prepontine region. **a.** Distribution of spinal projection neurons in the prepontine region. **b.** Confocal image series through the prepontine region from ventral (bottom) to dorsal (top). Each image represents a maximal projection stack of 20 µm. Proposed homology assignments for zebrafish are indicated in magenta dash lines. PnO, pontis oralis; LC, locus coeruleus; PB, parabrachial nucleus; PTg; LDT, Laterodorsal tegmentum; SR, superior raphe nucleus; mRT, midbrain reticular formation; CuF, cuneiform nucleus. **c.** Confocal images of the prepontine region showing the projecting neurons in the caudal SR after bilateral optical spinal backfills.

The pontine (P) region is defined rostrally by r2, containing the rostral trigeminal motor nucleus (5Nr), and caudally by rhombomere 4 (r4), which includes the Mauthner cell (M cell). In mammals, the bulk of spinal-projecting neurons in the pontine region is located in the pontis oralis nucleus of the reticular formation (PnO) (Liang et al., 2015, 2011a). This region is not well subdivided in mammals and contains spinal-projecting neurons with relatively large cell size (Liang et al., 2015). We found the same anatomical pattern in larval zebrafish, with spinal-projecting neurons with large somas located in the ventral planes, likely belonging to the PnO (**Figure 7c**). Dorsocaudally to the trigeminal motor nucleus, we found intensely labeled projecting neurons (**Figure 6b**, **Figure 7c**) likely belonging to the sensory trigeminal nuclei (s5) (Liang et al., 2015, 2011a). Caudal to the nuclei s5 at the ventrolateral alar plate in r4, the intermediate octavomotor nucleus (ION) shows numerous strongly labeled projecting neurons ipsilateral to the photoactivation site (**Figure 7c**) (Bussières et al., 1999).

**Figure 7.**
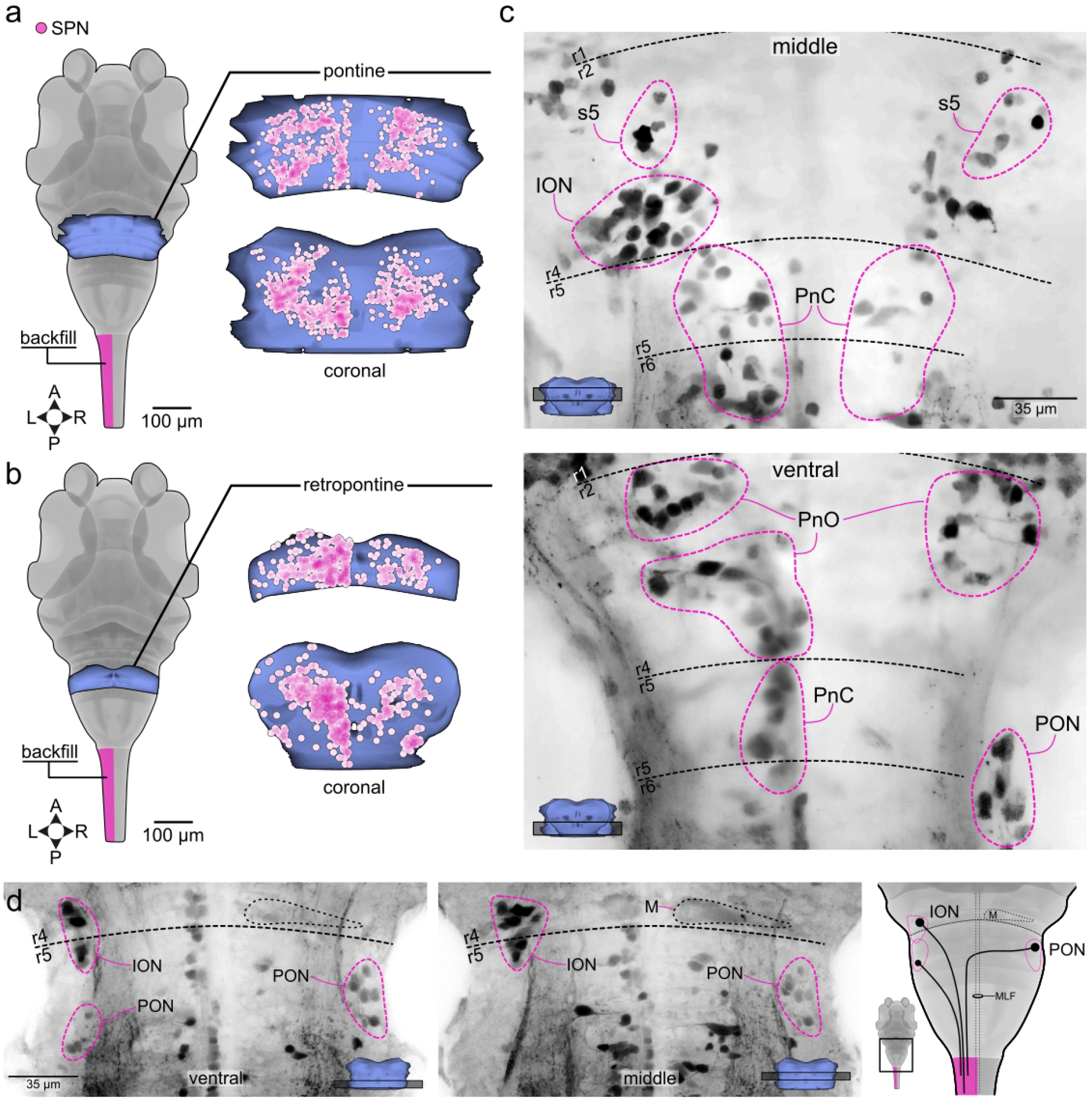
Rhombencephalic spinal-projecting neurons in the pontine and retropontine regions. **a.** Distribution of spinal projection neurons in the pontine region. **b.** Distribution of spinal projection neurons in the retropontine region. **c.** Confocal image series through the pontine-retropontine region from ventral (bottom) to dorsal (top). Each image represents a maximal projection stack of 20 µm. Proposed homology assignments for zebrafish are indicated in magenta dash lines. PnO, pontis oralis; PnC, pontis caudalis. ION, intermediate octavomotor nuclei; PON, posterior octavomotor nuclei; s5, sensory trigeminal nuclei. **d.** Confocal images of the pontine-retropontine region showing the distribution of the projecting neurons from the vestibular system in larval zebrafish. Each image represents a maximal projection stack of 10 µm. M, Mauthner cell.

The retropontine (RP) region consists of rhombomere 5 (r5) and rhombomere 6 (r6) containing the abducens (6N) and facial nucleus (7N), respectively. At this level, we found large and well-labeled neurons that belong to the pontine caudalis nucleus (PnC) (**Figure 7b, 7c**). Apart from the reticulospinal neurons, our backfill method in larval zebrafish also revealed the vestibulospinal system. In addition to the vestibulospinal neurons of the intermediate octavomotor nucleus (ION), we found that the posterior octavomotor nucleus (PON) contained both ipsilateral and contralateral spinal-projecting neurons (**Figure 7c-d**). The ipsilateral vestibulospinal neurons of the PON are located caudally and smaller than the contralateral ones (**Figure 7d**) (Bussières et al., 1999).

The medulla is rostrally delimited by rhombomere 7 (r7) and caudally by the rostral spinal cord (Watson et al., 2019b). Medially, it houses the gigantocellular reticular formation, a structure comprising multiple nuclei: the gigantocellular reticular nucleus (Gi), the alpha part of the gigantocellular reticular nucleus (GiA), the ventral part of the gigantocellular reticular nucleus (GiV), and the lateral paragigantocellular reticular nucleus (LPGi) (Liang et al., 2016, 2011b). Within the Gi, ipsilateral descending neurons are concentrated in the ventral region, while contralateral spinal-projecting neurons are prominent at its rostral pole (Liang et al., 2011b). Ventral to the Gi, the GiA, GiV, and LPGi all contain substantial numbers of spinal-projecting neurons with a strong ipsilateral tendency (Liang et al., 2016, 2011b). In mammals, the GiA/V and LPGi are ventrally located nuclei running alongside an extraraphe serotonergic column (VanderHorst and Ulfhake, 2006).

Our results show that in zebrafish, spinal-projecting neurons of the medulla display a slight ipsilateral predominance **(Figure 8a**, **Figure 1d**, **Figure 2c)**. Ventrally, the medially located inferior raphe nucleus (IR) is flanked by two columns of spinal-projecting neurons **(Figure 8b, c)**, which most likely correspond to the GiA/V or LPGi, given their colocalization with serotonergic neurons as labeled in the *Tg(pet1:GFP)* transgenic line **(Figure 8ci)**. Dorsal to these, a mass of densely packed spinal-projecting neurons with a sharp rostral boundary at r7 projects predominantly ipsilaterally to the spinal cord, and likely corresponds to the mammalian Gi (**Figure 8b, c**). Laterally, two well-defined bilaterally projecting columns likely correspond to the intermediate reticular nucleus (lRt), which separates the Gi from the more laterally situated parvocellular reticular nucleus (PCRt) (**Figure 8b, c**). At the caudal medulla, spinal-projecting neurons are more densely packed, making it challenging to distinguish putative homologous nuclei. Nevertheless, we propose that the ventral medullary reticular nucleus (MdV) lies caudal to the Gi, while the dorsal medullary reticular nucleus (MdD) is positioned lateral to the caudal pole of the lRt (**Figure 8b, c**). Finally, the spinal trigeminal nucleus (Sp5) appears to be delimited by well-labeled bilaterally projecting neurons located at the medullary border (**Figure 8c**).

**Figure 8.**
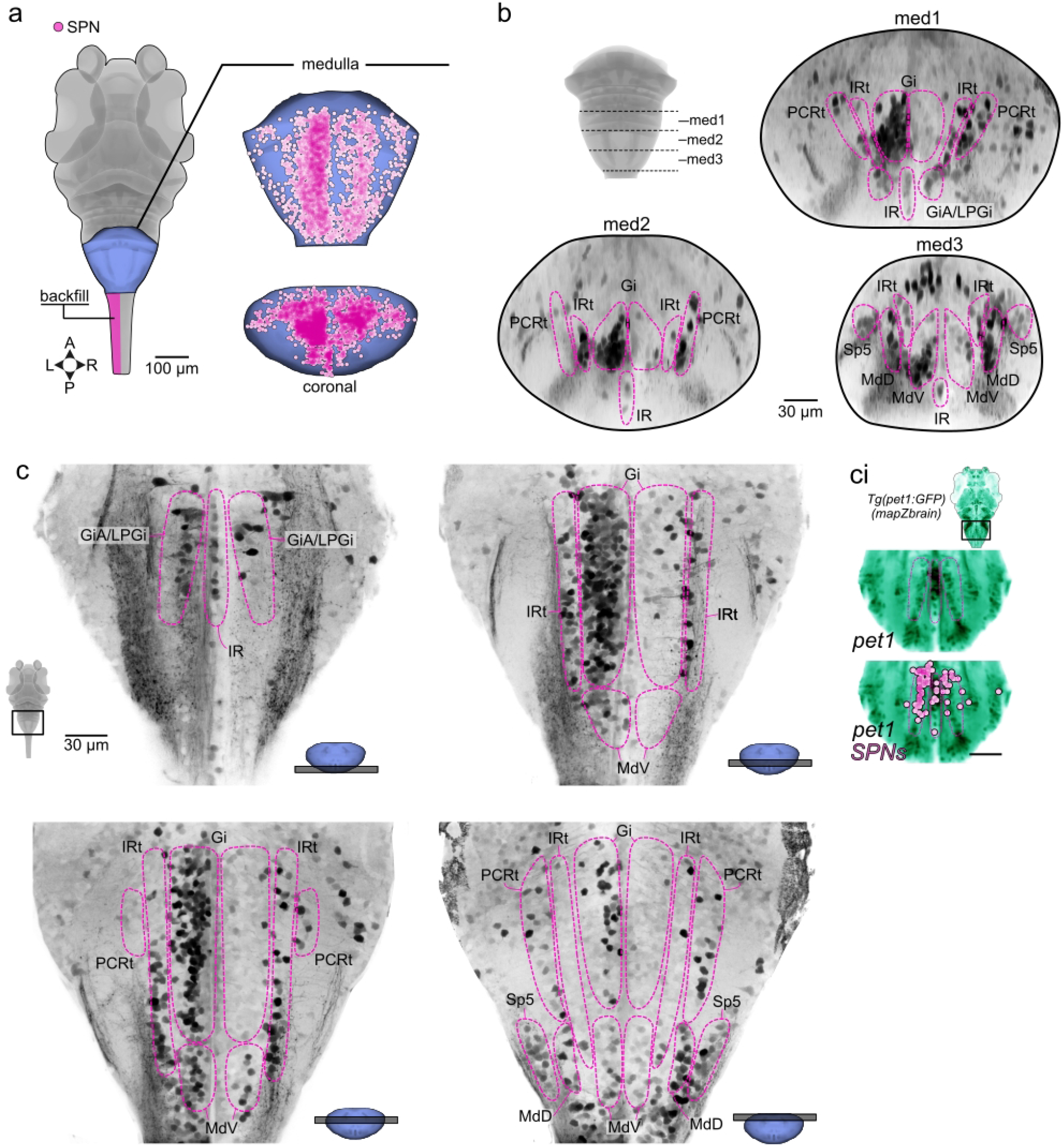
Rhombencephalic spinal-projecting neurons: Medullary region. **a.** Distribution of spinal projection neurons in the medullary region. **b.** Confocal image series in the coronal plane showing the spatial distribution of descending spinal projection neurons throughout the medulla from rostral (med1) to caudal (med3). **c.** Confocal image series through the medulla region from ventral (top-left) to dorsal (bottom-right). Each image represents a maximal projection stack of 20 µm. **ci.** Anatomical comparison of medullary GFP^+^ neurons in the *Tg(pet1:GFP)* line from the mapZbrain atlas with the location of the SPNs. Proposed homology assignments for zebrafish are indicated in magenta dash lines. GiA, gigantocellular reticular nucleus pars alpha; Gi, gigantocellular reticular nucleus; lRt, intermediate reticular nucleus; PCRt, parvocellular reticular nucleus; LPGi, lateral paragigantocellular nucleus; IR, inferior Raphe nucleus. MdV, ventral medullary reticular nucleus; MdD, dorsal medullary reticular nucleus. Sp5, spinal trigeminal nucleus.

## Discussion

This study provides a comprehensive description of the descending supraspinal system of larval zebrafish. Despite the proven track record of larval zebrafish as a powerful vertebrate model for systems neuroscience, descriptions of brain nuclei that project to the spinal cord have been surprisingly incomplete. Our approach revealed a complex descending system with an organization that rivals the one described in mammals (Cruce and Newman, 1984; Liang et al., 2011b; Nudo and Masterton, 1989; Tohyama et al., 1979). Using optical backfilling combined with two-photon photoactivation of PAGFP in transgenic larval zebrafish, we built the first comprehensive whole-brain map of spinal-projecting neurons (SPNs), identifying approximately ∼1,300 SPNs per larva distributed across all major brain regions. The hindbrain, particularly the medulla oblongata, contains the vast majority of SPNs, with the caudal reticular formation (Gi, GiA/V, LPGi) constituting the largest population. Strikingly, we detected telencephalic spinal projections originating from the olfactory bulb and subpallium, confirming that this pathway represents a conserved vertebrate feature rather than a mammalian innovation. In the diencephalon, distinct neuromodulatory populations were identified as SPNs, including oxytocinergic neurons in the preoptic area, dopaminergic neurons in the posterior tuberculum, and neurons of the accessory optic system. A key highlight of our work is the anatomical clarification of the interstitial pretectal nuclei: we propose that the GnRH2^+^ cluster corresponds to the Edinger-Westphal nucleus (EW), supported by co-expression of cart2 and uts1. In the mesencephalon, the red nucleus was identified as a compact cluster of just 5 neurons projecting contralaterally, and its connectivity with the cerebellum was confirmed using single-cell atlas data. Throughout the brainstem, our study identifies putative zebrafish homologs of key mammalian reticulospinal, vestibulospinal, and raphe-spinal populations, anchored by transgenic line expression data and brain atlas registration. Taken together, our findings offer a detailed description of the descending system in larval zebrafish that we hope will serve as a useful reference for both comparative vertebrate neuroanatomy and future functional work. Our findings indicate that the perceived ‘simplicity’ of the larval zebrafish brain reflects the constraints of classical labeling methods rather than an intrinsic biological property.

### Achievements and limitations of the study

The incomplete description of the zebrafish descending system reflects the limitations of earlier labeling methods. Retrograde tracing relies on tracer uptake through the axon, and older techniques preferentially labeled large, myelinated axons while missing smaller neurons with unmyelinated fibers (Beresford, 1965; Papez, 1926; Torvik and Brodal, 1957). While tracers like horseradish peroxidase (HRP) and small fluorescent dyes can allow visualization of finer reticulospinal projections (Cabot et al., 1982; Sánchez-Camacho et al., 2001; Ten Donkelaar, 1976), the original zebrafish description (Kimmel, 1982; Kimmel et al., 1982), performed alongside similar HRP backfills in other vertebrates, still yielded a low neuron count that made comparative analysis difficult (Cruce and Newman, 1984). This likely reflects a combination of factors: the early developmental stage of the larvae, the effectiveness of the mechanical damage caused by pressure-injecting dye into the spinal cord, and the possibility that small larval axons close too quickly for effective dye incorporation. In contrast, optical backfilling with two-photon imaging avoids many of these issues by allowing controlled axon labeling. The method does, however, have limitations. In particular, somatic signal intensity depends on the ratio of photoactivated PAGFP ascending from the axon to the volume of the cell body. Neurons with large somata projecting to spinal cord remain faint, which is why the Mauthner cells, despite having the largest axons, were consistently the faintest, a pattern also seen in large neurons of the INC, ND, and ventral SPNs at the PnO and PnC. Similarly, long-range neurons in the rostral diencephalon and telencephalon project thin axons to the spinal cord, resulting in faint somatic labeling and likely, underrepresentation in our data.

Traditionally, nuclei of supraspinal neurons projecting to spinal cord have been identified by first delineating brain regions using cytoarchitectonic criteria, and then confirming spinal projections through retrograde tracing (Cruce et al., 1999). Here, we took the opposite approach. Based on observations that the well-established principle that the vertebrate descending system is highly conserved across vertebrate species (Cruce et al., 1999; Cruce and Newman, 1984; Yamamoto et al., 2017); (Adli et al., 1999; Cabot et al., 1982; Cruce and Newman, 1981; Crutcher et al., 1978; Hermann et al., 2003; Sánchez-Camacho et al., 2001; Smeets and Timerick, 1981a; Ten Donkelaar, 1976; ten Donkelaar et al., 1980a, 1981a; Ten Donkelaar, 1982a), we used the location and projection patterns of retrogradely labeled neurons as an anchor for defining brain areas homology. This was a practical necessity, as the cytoarchitectonic boundaries in the brain of larval zebrafish are not as well defined as in other vertebrates, making anatomy alone an insufficient basis for homology comparisons.

The brain of 6-dpf larvae is actively developing, which could complicate comparisons with adult vertebrates. However, evidence from fish and other vertebrates suggests that the broad organization of the supraspinal descending system is established early in development (Auclair et al., 1993; Bosch and Roberts, 1994; Gross and Oppenheim, 1985), allowing us to make confident comparisons even at this developmental stage. Undoubtedly, the compact brain of a 6-dpf larva presents challenges, as many distinct nuclei are closely packed, making it difficult to delineate clear boundaries between adjacent groups. Whenever anatomical criteria were insufficient, we relied on molecular markers from the brain atlases to aid our comparative assessments. Altogether the homologies we proposed here should be considered as working hypotheses to be further tested using molecular developmental approaches to confirm homologies. We expect future studies integrating refined molecular, genetic, and functional data to challenge, expand, and refine the identifications proposed here (Pompeiano, 1973).

### Telencephalic descending projections

Comparative studies across vertebrates have consistently described a broadly conserved organization of descending nuclei and spinal pathways (Cruce et al., 1999; Cruce and Newman, 1984). The most rostrally located area consistently found to have descending projections is the diencephalic preoptic area (Cruce and Newman, 1981; New et al., 1998; Prasada Rao et al., 1987b; Smeets and Timerick, 1981b; Ten Donkelaar, 1982a, 1976; ten Donkelaar et al., 1981b, 1980a). Historically, no descending cells had been observed in the telencephalon of non-mammalian vertebrates (Ronan and Northcutt, 1985; ten Donkelaar, 1988; Ten Donkelaar, 1982a), with the exception of studies in nurse sharks, tiger salamander, iguana, and lamprey that found telencephalic projections reaching the rostral spinal cord using anterograde labeling (Ebbesson and Schroeder, 1971; Kokoros and Northcutt, 1977; Ocaña et al., 2015; Ten Donkelaar, 1982a). Our optical backfilling here revealed a few neurons with descending projections from the telencephalon in larval zebrafish. Later examination of the ZExplorer larval zebrafish brain and spinal cord atlas (Du et al., 2025) corroborated these sporadic observations. The morphology of the neurons in the atlas shows that these few telencephalic cells only project to the most rostral levels of the spinal cord. This explains why they are so difficult to capture retrogradely. Our findings challenge the established assumption that the telencephalic projections to the spinal cord are a derived feature of mammals (Cruce and Newman, 1981; New et al., 1998; Prasada Rao et al., 1987b; Smeets and Timerick, 1981b; Ten Donkelaar, 1982a, 1976; ten Donkelaar et al., 1981b, 1980a), and go in line with recent results from lampreys showing that the telencephalic (cortical) descending pathways appeared very early in vertebrate evolution (Ocaña et al., 2015).

### Diencephalic descending projections

The location and laterality of descending neurons in the hypothalamic preoptic area, rostral and intermediate hypothalamus, thalamus, and pretectum are consistent with descriptions in other early-diverging vertebrates, including tetrapods (Cruce and Newman, 1981; New et al., 1998; Prasada Rao et al., 1987b; Smeets and Timerick, 1981b; Ten Donkelaar, 1982a, 1976; ten Donkelaar et al., 1981b, 1980a). Our optical backfill results, combined by the expression patterns of various transgenic lines, corroborate that oxytocinergic and dopaminergic spinal inputs represent strongly conserved descending pathways across vertebrate species (Bohic et al., 2026; Oti et al., 2021; Ryczko et al., 2016; Sharples et al., 2014; Teclemariam-Mesbah et al., 1997). We paid particular attention to the delineation of the pretectal interstitial nuclei. Building on the premise that *gnrh2^+^* neurons correspond to the EW nucleus (Kauffman, 2004), we were able to more precisely define the nuclei situated lateral to EW. Across many vertebrate species, EW neurons express a wide range of peptides (Priest et al., 2023) regulating many behavioural aspects (Kauffman, 2004; Kozicz et al., 2011; Priest et al., 2023). Anatomically, the mammalian EW nucleus has wide range projections through the brain profusely projections to the spinal cord (Topilko et al., 2022; Yamamoto et al., 1995b). Besides a role in anxiety and depression-like behavior, mammalian EW-GnRH2 neurons are involved in regulating food intake (Kauffman, 2004; Kozicz et al., 2011). Similarly, in fish, modulation of GnRH2 signalling alters feeding behavior (Hofmann, 2006; Marvel et al., 2021, 2019; Nishiguchi et al., 2012), demonstrating a conserved functional role.

In mammals, the INC and ND are two interstitial nuclei that share considerable anatomical and functional similarities (Fukushima, 1991, 1987; Zhao et al., 2024). Both nuclei belong to the accessory oculomotor complex, classically implicated in the control of eye and neck movements (Fukushima, 1991, 1987; Zhao et al., 2024). A key anatomical criterion for distinguishing INC from ND is their relative position with respect to the MLF axons. We used this landmark to delineate these two nuclei as distinct clusters on either side of the MLF. In zebrafish, the INC and ND have historically been studied as a single entity under the designation of the nMLF (Kimmel et al., 1982). However, a recent study in larval zebrafish demonstrated that the nMLF can be subdivided into lateral and medial clusters based on differential expression of VGLUT2 and VGLUT1, respectively (Berg et al., 2023). This molecular segregation is further supported by physiological evidence, as the two clusters serve distinct functions in locomotor control. Such functional specialisation aligns well with our anatomical delineation of the INC and ND based on anatomical evidence. Similar to what we show here in larval zebrafish, the INC-ND sends descending projections to the caudal medulla in cats and mice (Skinner et al., 1984; Zhao et al., 2024). The role of the INC-ND in controlling locomotion (Berg et al., 2023; Kashin et al., 1974; Severi et al., 2014; Thiele et al., 2014) is not exclusive to zebrafish as electrical stimulations of this area shows locomotor promoting effects in cats and rats (Berezovskii, 1992; Berezovskiĭ, 1991; K.V.Bayev et al., 1986). Moreover, studies in rats showed that INC-ND is necessary for the execution of locomotion upon electrical stimulation of the hypothalamic locomotor region (Sinnamon and Benaur, 1997), further supporting the evolutionary conservation of the pretectal interstitial nuclei role in locomotion.

The location of the putative PAG in larval zebrafish was proposed solely on the basis of *relaxin 3a* (*rln3a*) gene expression (Donizetti et al., 2009, 2008; Spikol et al., 2024), without consideration of anatomical landmarks or comparative neuroanatomical criteria (Donizetti et al., 2009, 2008; Spikol et al., 2024). A recent report indicates that the neurons located in the above region tentatively labeled as PAG send ipsilateral descending spinal projections (Spikol et al., 2024). We challenge this designation of PAG in larval zebrafish. In mammals, RLN3 expression within the PAG is sparse and largely confined to its dorsal subdivision (Smith et al., 2010), a region devoid of spinal-projecting neurons (Liang et al., 2011b). By contrast, RLN3 is far more prominently expressed in the mRT, located laterally to the RN, with clear descending projections to the spinal cord (Liang et al., 2011b; Smith et al., 2010). The population described by Spikol and collaborators (Spikol et al., 2024) aligns anatomically with the characteristics of the mRT rather than the PAG, and we therefore propose that their designation represents a misidentification of these two structures. Instead, we propose that the PAG in larval zebrafish is located medially and dorsal to the interstitial nuclei INC, ND, and EW, a position that better respects its defining anatomical relationship to the cerebral aqueduct and is consistent with the location described across vertebrates (Olson et al., 2017). It is worth noting that as development progresses, the expansion of the cerebral aqueduct displaces the PAG away from the midline, creating the impression of a more lateral position. However, its relative location with respect to surrounding nuclei remains unchanged throughout (Goodson and Bass, 2002; Kittelberger and Bass, 2013)

### Mesencephalic pathways

The tectospinal neurons were not not identified in teleosts (Kimmel et al., 1982; Lee and Eaton, 1991b; Oka et al., 1986; Prasada Rao et al., 1987b), but found in elasmobranchs (Smeets and Timerick, 1981a) and lungfishes (Ronan and Northcutt, 1985). The descending tectal afferent neurons reach only the most rostral spinal levels (Altman and Carpenter, 1961; Foster and Hall, 1975; Luiten, 1981; Rubinson, 2008). In contrast to previous reports (New et al., 1998; Oka et al., 1986; Prasada Rao et al., 1987b), our approach revealed, that actinopterygian fishes possess tectospinal projections matching the locations and laterality described for other vertebrate species. The red nucleus (RN) is defined by its relative position in the tegmentum mesencephali, at the level of the oculomotor nerve, rostral to the decussation of the brachium conjunctivum. Classically, the criteria used to identify a structure as RN are, apart from its relative position, its contralateral spinal projection (ten Donkelaar, 1988). Here, we identified the RN in larval zebrafish as a small and compact cluster of neurons located in the tegmentum, lateral to the oculomotor nucleus, with clear contralateral spinal projections. In teleosts, the contralaterally projecting RN is defined as the “nucleus ruber of Goldstein” (NRg) (Goldstein, 1905; Prasada Rao et al., 1987b; Yamamoto et al., 2017).

### Reticulospinal neurons: conserved core of motor control

The reticular formation is a phylogenetically conserved collection of neurons dispersed throughout the medulla, pons, and isthmus, embedded within a dense network of longitudinally and commissurally projecting fibers (Balcells, n.d.; Brodal, 1957; Pompeiano, 1973). While Meessen and Olszewski (Meessen and Olszewski, 1949) established the modern mammalian nomenclature of the reticular formation based on cytoarchitectonically distinct nuclei, this precise parcellation cannot be directly extended to non-mammalian vertebrate species, for which the reticular formation has not been characterized with the same anatomical resolution (Cruce and Nieuwenhuys, 1974; Nieuwenhuys, 1974, 1972). Reticulospinal neurons constitute the most ancestral descending system involved in motor control in all classes of vertebrates (ten Donkelaar, 1988). By using the location and projection patterns of retrogradely labeled neurons as criteria for defining tentative homologous brain areas, we were able to confidently assign designations to the reticular formation of larval zebrafish, relying on comparisons with mammalian nuclear identification.

To place our findings in a broader evolutionary context, we compiled and compared data on reticulospinal neurons across vertebrate species (**Figure 9**). For this comparison, we excluded both vestibulospinal neurons, which fall outside the reticular formation proper (Brodal, 1957; Pompeiano, 1973; Ten Donkelaar, 1982a), and raphespinal neurons, whose representation is inconsistent across specimens due to their unreliable labeling with classical tracing techniques. To further facilitate cross-species comparisons, we follow the parcellation proposed by Hoevell (Hoevell, J.J.L.D. van, 1911), who divided the reticular formation into three regions based on the arrangement of its largest cells: the superior reticular nucleus (SRN), spanning from the isthmus to the rostral trigeminal nucleus in rhombomere 2; the middle reticular nucleus (MRN), extending from rhombomere 3 to rhombomere 6 at the level of the facial nucleus; and the inferior reticular nucleus (IRN), running from rhombomere 7 to the obex. We also follow the convention of excluding midbrain and diencephalic structures from the reticular formation (Brodal, 1957; Pompeiano, 1973). This comparative analysis reveals a striking degree of conservation in the topographical organization of reticulospinal neurons across vertebrates. In the IRN, these neurons are densely packed and medially concentrated, with their distribution terminating abruptly at rhombomere 6. Within the MRN and SRN, reticulospinal neurons fan out into two distinct extensions toward the isthmic region, giving the overall distribution a characteristic Y-shape in dorsal view (**Figure 9**). Our zebrafish results fit seamlessly within this conserved organizational plan, reinforcing the view that the fundamental architecture of the reticulospinal system was established early in vertebrate evolution (ten Donkelaar, 2009). Within this conserved structural plan, early-diverging vertebrate species tend to have fewer but larger reticulospinal neurons compared to mammals (Cruce and Nieuwenhuys, 1974; Heijdra and Nieuwenhuys, 1994; Newman and Cruce, 1982; Nieuwenhuys, 2011; Nieuwenhuys and Oey, 1983; Opdam et al., 1976), and the progressive increase in neuron number across lineages likely reflects the growing complexity of the body plan, the emergence of appendages, and the accompanying expansion of the motor repertoire.

**Figure 9.**
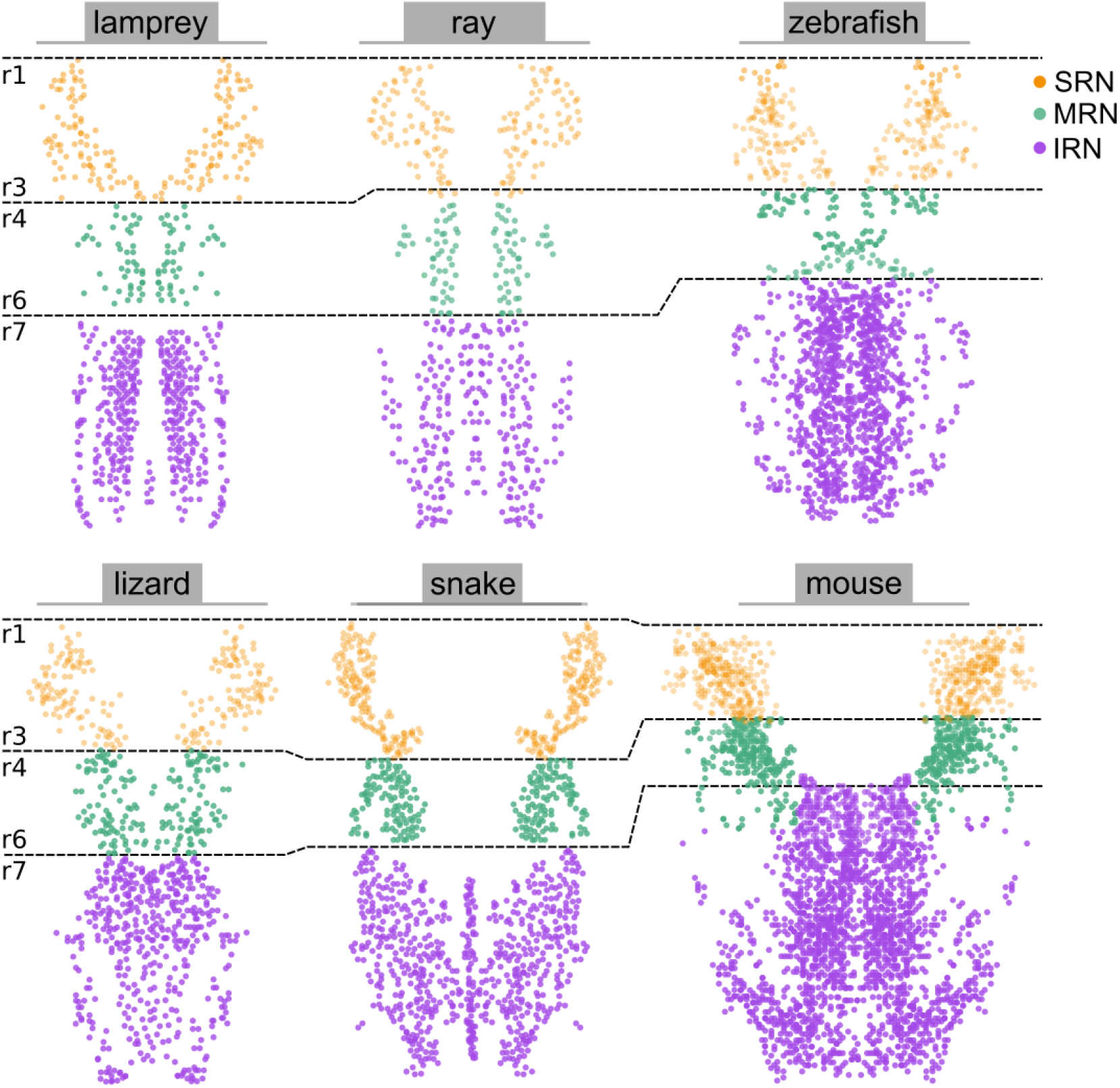
Comparative distribution of reticulospinal neurons across vertebrates. Dorsal view schematic illustrating the distribution of reticulospinal neurons in six vertebrate species: lamprey larvae, ray, larval zebrafish, lizard, snake, and mouse. Each dot represents a single retrogradely labeled reticulospinal neurons. Neurons are color-coded according to their assignment to one of three reticular subdivisions following the parcellation of Hoevell (1911): the superior reticular nucleus (SRN, orange), spanning from the isthmus to rhombomere 2; the middle reticular nucleus (MRN, green), extending from rhombomere 3 to rhombomere 6; and the inferior reticular nucleus (IRN, purple), running from rhombomere 7 to the obex. Dashed lines indicate rhombomere boundaries. Rostral is up. Data for each species were compiled from: lamprey larvae (Swain et al., 1993), ray (Smeets and Timerick, 1981b), lizard (Wolters et al., 1982), snake (Ten Donkelaar, 1982a), and mouse (Wang et al., 2022). Vestibulospinal and raphespinal neurons were excluded from this comparison (see text).

## Contributions

M.C.T: Conceptualization, methodology, formal investigation, formal analysis, plotting, literature review, writing original draft, review and editing. K.F: Reviewing and editing. T.W: Assisted performing initial backfills experiments. S.N, and M.A. Generated Tg(elavl3:stGtACR1-FusionRed) transgenic line. R.D: Review and editing. C.W: Conceptualization, review and editing, funding acquisition.

## Data availability

Data and code supporting the findings of this study are publicly available on the Open Science Framework (https://osf.io/). The repository includes the source coordinate dataset and R analysis script. The repository can be accessed at https://doi.org/10.17605/OSF.IO/C6F4Z.

## Acknowledgements

We deeply thank Professor Yonathan Zohar for sharing the Tg*(GnRH2:EGFP)* transgenic line. We thank Sophie Nunes Figueiredo, Camille Lejeune, Antoine Arneau and Nolwenn Jezequel From Paris Brain Institute PHENO Zfish Core Facility, RRID:SCR_028198. We thank Dr. Olivier Renaud, Claire Lovo from the ICM.Quant core facility.

## References

Adli DSH, Stuesse SL, Cruce WLR. 1999. Immunohistochemistry and spinal projections of the reticular formation in the northern leopard frog, Rana pipiens. Journal of Comparative Neurology 404:387–407. DOI: 10.1002/(SICI)1096-9861(19990215)404:3%3C387::AID-CNE8%3E3.0.CO;2-Z

Agetsuma M, Aizawa H, Aoki T, Nakayama R, Takahoko M, Goto M, Sassa T, Amo R, Shiraki T, Kawakami K, Hosoya T, Higashijima S, Okamoto H. 2010. The habenula is crucial for experience-dependent modification of fear responses in zebrafish. Nature Neuroscience 13:1354–1356. DOI: 10.1038/nn.2654

Altman J, Carpenter MB. 1961. Fiber projections of the superior colliculus in the cat. Journal of Comparative Neurology 116:157–177. DOI: 10.1002/cne.901160206

Ando R, Hama H, Yamamoto-Hino M, Mizuno H, Miyawaki A. 2002. An optical marker based on the UV-induced green-to-red photoconversion of a fluorescent protein. Proceedings of the National Academy of Sciences 99:12651–12656. DOI: 10.1073/pnas.202320599

Auclair F, Bélanger MC, Marchand R. 1993. Ontogenetic study of early brain stem projections to the spinal cord in the rat. Brain Research Bulletin 30:281–289. DOI: 10.1016/0361-9230(93)90256-b, PMID: 8457877

Balcells M. n.d. Historical aspects of the anatomy of the reticular formation.

Beresford WA. 1965. A Discussion on Retrograde Changes in Nerve Fibres. In: Singer M, Schadé JP (Eds). Progress in Brain Research, Degeneration Patterns in the Nervous System. Elsevier. p. 33–56. DOI: 10.1016/S0079-6123(08)63738-3

Berezovskii VK. 1992. Brainstem pathways of the initiation of locomotion. Neurophysiology 23:357–371. DOI: 10.1007/BF01052569

Berezovskiĭ VK. 1991. [The participation of the interstitial nucleus of Cajal in initiating locomotion in cats and rats]. Neirofiziologiia = Neurophysiology 23:368–371. PMID: 1881493

Berg EM, Mrowka L, Bertuzzi M, Madrid D, Picton LD, Manira AE. 2023. Brainstem circuits encoding start, speed, and duration of swimming in adult zebrafish. Neuron 111:372–386.e4. DOI: 10.1016/j.neuron.2022.10.034, PMID: 36413988

Bhandiwad AA, Chu NC, Semenova SA, Holmes GA, Burgess HA. 2022. A cerebellar-prepontine circuit for tonic immobility triggered by an inescapable threat. Science Advances 8:eabo0549. DOI: 10.1126/sciadv.abo0549

Bhandiwad AA, Gupta T, Subedi A, Heigh V, Holmes GA, Burgess HA. 2024. Brain Imaging and Registration in Larval Zebrafish. In: Amatruda JF, Houart C, Kawakami K, Poss KD (Eds). Zebrafish: Methods and Protocols. Springer US. p. 141–153. DOI: 10.1007/978-1-0716-3401-1_9

Bianco IH, Ma L-H, Schoppik D, Robson DN, Orger MB, Beck JC, Li JM, Schier AF, Engert F, Baker R. 2012. The tangential nucleus controls a gravito-inertial vestibulo-ocular reflex. Current biology: CB 22:1285–1295. DOI: 10.1016/j.cub.2012.05.026, PMID: 22704987

Bluett RJ, Yu Y, Pauli JL, Campos CA, Palmiter RD. 2025. Edinger-Westphal Urocortin-1 neurons regulate consumption and affect. Cell Reports 44. DOI: 10.1016/j.celrep.2025.115814, PMID: 40512623

Bohic M, Salamone PC, Zuo W, Negm A, Fulton SL, Du S, Jayakumar S, Keating J, Soubeyre V, Gradwell MA, Upadhyay A, Shorter L, Kim J, Inoue YU, Inoue T, Mensch B, Ye J-H, Peirs C, Poulen G, Lonjon N, Vachiery-Lahaye F, Bauchet L, Bourinet E, Olausson H, Abdus-Saboor I, Tao Y-X, Boehme R, Abraira VE. 2026. Oxytocin Modulation of Spinal Circuits Drives Therapeutic Benefits of Massage. DOI: 10.64898/2026.01.11.698886

Bosch TJ, Roberts BL. 2001. The Relationships of Brain Stem Systems to Their Targets in the Spinal Cord of the Eel, Anguilla anguilla. Brain Behavior and Evolution 57:106–116. DOI: 10.1159/000047230

Bosch TJ, Roberts BL. 1994. The Size and Number of Neurons Descending to the Spinal Cord in Relation to Body Length in the European Eel (Anguilla anguilla). Brain Behavior and Evolution 44:50–60. DOI: 10.1159/000113569

Brodal A. 1957. The reticular formation of the brain stem. Anatomical aspects and functionnal correlations. The Henderson Trust Lectures 87.

Burgess HA, Burton EA. 2023. A Critical Review of Zebrafish Neurological Disease Models−1. The Premise: Neuroanatomical, Cellular and Genetic Homology and Experimental Tractability. Oxford Open Neuroscience 2:kvac018. DOI: 10.1093/oons/kvac018, PMID: 37649777

Bussières N, Pflieger J-F, Dubuc R. 1999. Anatomical study of vestibulospinal neurons in lampreys. Journal of Comparative Neurology 407:512–526. DOI: 10.1002/(SICI)1096-9861(19990517)407:4%3C512::AID-CNE4%3E3.0.CO;2-%2523

Cabot JB, Reiner A, Bogan N. 1982. Avian bulbospinal pathways: anterograde and retrograde studies of cells of origin, funicular trajectories and laminar terminations. Progress in Brain Research 57:79–108. DOI: 10.1016/S0079-6123(08)64125-4, PMID: 6296922

Carbo-Tano M, Lapoix M, Jia X, Thouvenin O, Pascucci M, Auclair F, Quan FB, Albadri S, Aguda V, Farouj Y, Hillman EMC, Portugues R, Del Bene F, Thiele TR, Dubuc R, Wyart C. 2023. The mesencephalic locomotor region recruits V2a reticulospinal neurons to drive forward locomotion in larval zebrafish. Nature Neuroscience 26:1775–1790. DOI: 10.1038/s41593-023-01418-0

Castiglioni AJ, Gallaway MC, Coulter JD. 1978. Spinal projections from the midbrain in monkey. The Journal of Comparative Neurology 178:329–346. DOI: 10.1002/cne.901780208, PMID: 415074

Collins EMD, Silva PTM, Ostrovsky AD, Renninger SL, Tomás AR, Corral RD del, Orger MB. 2025. Characterisation of transgenic lines labelling reticulospinal neurons in larval zebrafish. eNeuro. DOI: 10.1523/ENEURO.0581-24.2025, PMID: 40374558

Cruce WL, Newman DB. 1981. Brain stem origins of spinal projections in the lizard Tupinambis nigropunctatus. The Journal of Comparative Neurology 198:185–207. DOI: 10.1002/cne.901980202, PMID: 7240441

Cruce WL, Nieuwenhuys R. 1974. The cell masses in the brain stem of the turtle Testudo hermanni; a topographical and topological analysis. The Journal of Comparative Neurology 156:277–306. DOI: 10.1002/cne.901560303, PMID: 4418301

Cruce WLR, Newman DB. 1984. Evolution of Motor Systems: The Reticulospinal Pathways. American Zoologist 24:733–753. DOI: 10.1093/icb/24.3.733

Cruce WLR, Stuesse SL, Northcutt RG. 1999. Brainstem neurons with descending projections to the spinal cord of two elasmobranch fishes: Thornback guitarfish, Platyrhinoidis triseriata, and horn shark, Heterodontus francisci. Journal of Comparative Neurology 403:534–560. DOI: 10.1002/(SICI)1096-9861(19990125)403:4%3C534::AID-CNE8%3E3.0.CO;2-8

Crutcher KA, Humbertson AO, Martin GF. 1978. The origin of brainstem-spinal pathways in the North American opossum (Didelphis virginiana). Studies using the horseradish peroxidase method. The Journal of Comparative Neurology 179:169–193. DOI: 10.1002/cne.901790111, PMID: 8980723

Donizetti A, Fiengo M, Minucci S, Aniello F. 2009. Duplicated zebrafish relaxin-3 gene shows a different expression pattern from that of the co-orthologue gene. Development, Growth & Differentiation 51:715–722. DOI: 10.1111/j.1440-169X.2009.01131.x

Donizetti A, Grossi M, Pariante P, D’Aniello E, Izzo G, Minucci S, Aniello F. 2008. Two neuron clusters in the stem of postembryonic zebrafish brain specifically express relaxin-3 gene: first evidence of nucleus incertus in fish. Developmental Dynamics: An Official Publication of the American Association of Anatomists 237:3864–3869. DOI: 10.1002/dvdy.21786, PMID: 18985751

Du X-F, Yue Z-F, Chen M-Q, Li W-L, Chen T-L, Hu H-Y, Wan H-L, Zhao T-T, Zhong Y-W, Ning X-Y, Zheng X-D, Ren H-A, Wang R-Q, Zhao R, Peng X-L, Jia Z-M, Jin C-X, Huang J-W, Deng W, Qian L-Q, Wang L, Zheng M-Y, Zhang W, Shen X-Y, Shen X-L, Qiu X-Y, Zhao Q-M, Li D-Y, Chen L-J, Wang S-J, Gong Y-C, Li Y, Mu Y, Ji P, He J, Wang Y-F, Du J-L. 2025. Central nervous system atlas of larval zebrafish based on the morphology of single excitatory and inhibitory neurons. DOI: 10.1101/2025.06.06.658008

Ebbesson SO, Schroeder DM. 1971. Connections of the nurse shark’s telencephalon. Science 173:254–256. DOI: 10.1126/science.173.3993.254, PMID: 5087492

Fedorov A, Beichel R, Kalpathy-Cramer J, Finet J, Fillion-Robin J-C, Pujol S, Bauer C, Jennings D, Fennessy F, Sonka M, Buatti J, Aylward S, Miller JV, Pieper S, Kikinis R. 2012. 3D Slicer as an image computing platform for the Quantitative Imaging Network. Magnetic Resonance Imaging 30:1323–1341. DOI: 10.1016/j.mri.2012.05.001, PMID: 22770690

Finger TE. 1978. Efferent neurons of the teleost cerebellum. Brain Research 153:608–614. DOI: 10.1016/0006-8993(78)90346-3

Foster RE, Hall WC. 1975. The connections and laminar organization ofthe optic tectum in a reptile (lguana iguana). The Journal of Comparative Neurology 163:397–425. DOI: 10.1002/cne.901630403, PMID: 1176645

François A, Low SA, Sypek EI, Christensen AJ, Sotoudeh C, Beier KT, Ramakrishnan C, Ritola KD, Sharif-Naeini R, Deisseroth K, Delp SL, Malenka RC, Luo L, Hantman AW, Scherrer G. 2017. A Brainstem-Spinal Cord Inhibitory Circuit for Mechanical Pain Modulation by GABA and Enkephalins. Neuron 93:822–839.e6. DOI: 10.1016/j.neuron.2017.01.008

Fukushima K. 1991. The interstitial nucleus of Cajal in the midbrain reticular formation and vertical eye movement. Neuroscience Research 10:159–187. DOI: 10.1016/0168-0102(91)90055-4, PMID: 1650435

Fukushima K. 1987. The interstitial nucleus of Cajal and its role in the control of movements of head and eyes. Progress in Neurobiology 29:107–192. DOI: 10.1016/0301-0082(87)90016-5, PMID: 3108957

Gahtan E, O’Malley DM. 2003. Visually guided injection of identified reticulospinal neurons in zebrafish: a survey of spinal arborization patterns. The Journal of Comparative Neurology 459:186–200. DOI: 10.1002/cne.10621, PMID: 12640669

Goldstein K. 1905. Untersuchungen über das Vorderhirn und Zwischenhirn einiger Knochenfische (nebsteinigen Beiträgen über Mittelhirn und Kleinhirn derselben.). Archiv für mikroskopische Anatomie 66:135–219. DOI: 10.1007/BF02979208

Goodson JL, Bass AH. 2002. Vocal–acoustic circuitry and descending vocal pathways in teleost fish: Convergence with terrestrial vertebrates reveals conserved traits. Journal of Comparative Neurology 448:298–322. DOI: 10.1002/cne.10258

Gross GH, Oppenheim RW. 1985. Novel sources of descending input to the spinal cord of the hatchling chick. Journal of Comparative Neurology 232:162–179. DOI: 10.1002/cne.902320203

Heijdra YF, Nieuwenhuys R. 1994. Topological analysis of the brainstem of the bowfin, Amia calva. Journal of Comparative Neurology 339:12–26. DOI: 10.1002/cne.903390104

Hermann GE, Holmes GM, Rogers RC, Beattie MS, Bresnahan JC. 2003. Descending spinal projections from the rostral gigantocellular reticular nuclei complex. The Journal of Comparative Neurology 455:210–221. DOI: 10.1002/cne.10455, PMID: 12454986

Hlavacek M, Tahar M, Libouban S, Szabo T. 1984. The mormyrid brainstem. I. Distribution of brainstem neurones projecting to the spinal cord in Gnathonemus petersii. An HRP study. Journal Fur Hirnforschung 25:603–615. PMID: 6526990

Hoevell, J.J.L.D. van. 1911. Remarks on the reticular cells of the oblongata in different vertebrates. Proc. Acad. Sci. Amsterdam 13:1047–1065.

Hofmann HA. 2006. Gonadotropin-releasing hormone signaling in behavioral plasticity. Current Opinion in Neurobiology 16:343–350. DOI: 10.1016/j.conb.2006.05.005, PMID: 16697636

Huang K-H, Ahrens MB, Dunn TW, Engert F. 2013. Spinal Projection Neurons Control Turning Behaviors in Zebrafish. Current Biology 23:1566–1573. DOI: 10.1016/j.cub.2013.06.044, PMID: 23910662

Ingram WR, Ranson SW. 1935. THE NUCLEUS OF DARKSCHEWITSCH AND NUCLEUS INTERSTITIALIS IN THE BRAIN OF MAN. The Journal of Nervous and Mental Disease 81:125.

Kashin SM, Feldman AG, Orlovsky GN. 1974. Locomotion of fish evoked by electrical stimulation of the brain. Brain Research 82:41–47. DOI: 10.1016/0006-8993(74)90891-9, PMID: 4611595

Kauffman AS. 2004. Emerging functions of gonadotropin-releasing hormone II in mammalian physiology and behaviour. Journal of Neuroendocrinology 16:794–806. DOI: 10.1111/j.1365-2826.2004.01229.x, PMID: 15344918

Kim LH, Sharma S, Sharples SA, Mayr KA, Kwok CHT, Whelan PJ. 2017. Integration of Descending Command Systems for the Generation of Context-Specific Locomotor Behaviors. Frontiers in Neuroscience 11:581. DOI: 10.3389/fnins.2017.00581, PMID: 29093660

Kimmel CB. 1982. Reticulospinal and vestibulospinal neurons in the young larva of a teleost fish, Brachydanio rerio. Progress in Brain Research 57:1–23. DOI: 10.1016/S0079-6123(08)64122-9, PMID: 7156394

Kimmel CB, Powell SL, Metcalfe WK. 1982. Brain neurons which project to the spinal cord in young larvae of the zebrafish. The Journal of Comparative Neurology 205:112–127. DOI: 10.1002/cne.902050203, PMID: 7076887

Kimura Y, Okamura Y, Higashijima S. 2006. alx, a zebrafish homolog of Chx10, marks ipsilateral descending excitatory interneurons that participate in the regulation of spinal locomotor circuits. The Journal of Neuroscience: The Official Journal of the Society for Neuroscience 26:5684–5697. DOI: 10.1523/JNEUROSCI.4993-05.2006, PMID: 16723525

Kimura Y, Satou C, Fujioka S, Shoji W, Umeda K, Ishizuka T, Yawo H, Higashijima S. 2013. Hindbrain V2a Neurons in the Excitation of Spinal Locomotor Circuits during Zebrafish Swimming. Current Biology 23:843–849. DOI: 10.1016/j.cub.2013.03.066, PMID: 23623549

Kittelberger JM, Bass AH. 2013. Vocal-Motor and Auditory Connectivity of the Midbrain Periaqueductal Gray in a Teleost Fish. The Journal of comparative neurology 521:791–812. DOI: 10.1002/cne.23202, PMID: 22826153

Kokoros JJ, Northcutt RG. 1977. Telencephalic efferents of the tiger salamander Ambystoma tigrinurn tigrinum (Green). Journal of Comparative Neurology 173:613–627. DOI: 10.1002/cne.901730402

Kozicz T, Bittencourt JC, May PJ, Reiner A, Gamlin PDR, Palkovits M, Horn AKE, Toledo CAB, Ryabinin AE. 2011. THE EDINGER-WESTPHAL NUCLEUS: A HISTORICAL, STRUCTURAL AND FUNCTIONAL PERSPECTIVE ON A DICHOTOMOUS TERMINOLOGY. The Journal of comparative neurology 519:1413–1434. DOI: 10.1002/cne.22580, PMID: 21452224

Kunst M, Laurell E, Mokayes N, Kramer A, Kubo F, Fernandes AM, Förster D, Dal Maschio M, Baier H. 2019. A Cellular-Resolution Atlas of the Larval Zebrafish Brain. Neuron 103:21–38.e5. DOI: 10.1016/j.neuron.2019.04.034, PMID: 31147152

Kuypers HG, Maisky VA. 1975. Retrograde axonal transport of horseradish peroxidase from spinal cord to brain stem cell groups in the cat. Neuroscience Letters 1:9–14. DOI: 10.1016/0304-3940(75)90004-x, PMID: 19604744

Kuypers HGJM. 2011. Anatomy of the Descending Pathways. Comprehensive Physiology. John Wiley & Sons, Ltd. p. 597–666. DOI: 10.1002/cphy.cp010213

Kuypers HGJM. 1964. The Descending Pathways to the Spinal Cord, their Anatomy and Function. In: Eccles JC, Schadé JP (Eds). Progress in Brain Research, Organization of the Spinal Cord. Elsevier. p. 178–202. DOI: 10.1016/S0079-6123(08)64048-0

K.V. Bayev, Berezovskii V, V.B. Esipenko. 1986. New Locomotor Regions in the Brainstem of Cat. Neirofiziologiia = Neurophysiology 18:416–419.

Lacroix-Ouellette P, Dubuc R. 2023. Brainstem neural mechanisms controlling locomotion with special reference to basal vertebrates. Frontiers in Neural Circuits 17.

Lau JYN, Fitzgerald JE, Bianco IH. 2025. Supraspinal commands have a modular organization that is behavioral context specific. Current Biology 0. DOI: 10.1016/j.cub.2025.07.066, PMID: 40848723

Lee RK, Eaton RC. 1991a. Identifiable reticulospinal neurons of the adult zebrafish, Brachydanio rerio. The Journal of Comparative Neurology 304:34–52. DOI: 10.1002/cne.903040104, PMID: 2016411

Lee RK, Eaton RC. 1991b. Identifiable reticulospinal neurons of the adult zebrafish, Brachydanio rerio. The Journal of Comparative Neurology 304:34–52. DOI: 10.1002/cne.903040104, PMID: 2016411

Lee RK, Eaton RC, Zottoli SJ. 1993. Segmental arrangement of reticulospinal neurons in the goldfish hindbrain. The Journal of Comparative Neurology 329:539–556. DOI: 10.1002/cne.903290409, PMID: 8454739

Leiras R, Cregg JM, Kiehn O. 2022. Brainstem Circuits for Locomotion. Annual Review of Neuroscience 45:63–85. DOI: 10.1146/annurev-neuro-082321-025137

Liang H, Paxinos G, Watson C. 2011a. Projections from the brain to the spinal cord in the mouse. Brain Structure & Function 215:159–186. DOI: 10.1007/s00429-010-0281-x, PMID: 20936329

Liang H, Paxinos G, Watson C. 2011b. Projections from the brain to the spinal cord in the mouse. Brain Structure & Function 215:159–186. DOI: 10.1007/s00429-010-0281-x, PMID: 20936329

Liang H, Watson C, Paxinos G. 2016. Terminations of reticulospinal fibers originating from the gigantocellular reticular formation in the mouse spinal cord. Brain Structure & Function 221:1623–1633. DOI: 10.1007/s00429-015-0993-z, PMID: 25633472

Liang H, Watson C, Paxinos G. 2015. Projections from the oral pontine reticular nucleus to the spinal cord of the mouse. Neuroscience Letters 584:113–118. DOI: 10.1016/j.neulet.2014.10.025, PMID: 25459287

Liu Y, Latremoliere A, Li X, Zhang Z, Chen M, Wang X, Fang C, Zhu J, Alexandre C, Gao Z, Chen B, Ding X, Zhou J-Y, Zhang Y, Chen C, Wang KH, Woolf CJ, He Z. 2018. Touch and tactile neuropathic pain sensitivity are set by corticospinal projections. Nature 561:547–550. DOI: 10.1038/s41586-018-0515-2

Luiten PG. 1981. Afferent and efferent connections of the optic tectum in the carp (Cyprinus carpio L.). Brain Research 220:51–65. DOI: 10.1016/0006-8993(81)90210-9, PMID: 6168333

Marvel M, Levavi-Sivan B, Wong T-T, Zmora N, Zohar Y. 2021. Gnrh2 maintains reproduction in fasting zebrafish through dynamic neuronal projection changes and regulation of gonadotropin synthesis, oogenesis, and reproductive behaviors. Scientific Reports 11:6657. DOI: 10.1038/s41598-021-86018-3

Marvel MM, Spicer OS, Wong T-T, Zmora N, Zohar Y. 2019. Knockout of Gnrh2 in zebrafish (Danio rerio) reveals its roles in regulating feeding behavior and oocyte quality. General and Comparative Endocrinology 280:15–23. DOI: 10.1016/j.ygcen.2019.04.002, PMID: 30951724

Meessen H, Olszewski J. 1949. Cytoarchitektonischer Atlas des Rautenhirns des Kaninchens. DOI: 10.1159/000426870

New JG, Snyder BD, Woodson KL. 1998. Descending neural projections to the spinal cord in the channel catfish, Ictalurus punctatus. The Anatomical Record 252:235–253. DOI: 10.1002/(SICI)1097-0185(199810)252:2%3C235::AID-AR9%3E3.0.CO;2-1, PMID: 9776078

Newman DB, Cruce WLR. 1982. The organization of the reptilian brainstem reticular formation: A comparative study using Nissl and Golgi techniques. Journal of Morphology 173:325–349. DOI: 10.1002/jmor.1051730309

Nieuwenhuys R. 2011. The Structural, Functional, and Molecular Organization of the Brainstem. Frontiers in Neuroanatomy 5. DOI: 10.3389/fnana.2011.00033

Nieuwenhuys R. 1974. Topological analysis of the brain stem: A general introduction. Journal of Comparative Neurology 156:255–276. DOI: 10.1002/cne.901560302

Nieuwenhuys R. 1972. Topological analysis of the brain stem of the lamprey Lampetra fluviatilis. Journal of Comparative Neurology 145:165–177. DOI: 10.1002/cne.901450204

Nieuwenhuys R, Oey PL. 1983. Topological analysis of the brainstem of the reedfish, Erpetoichthys calabaricus. The Journal of Comparative Neurology 213:220–232. DOI: 10.1002/cne.902130208, PMID: 6841670

Nishiguchi R, Azuma M, Yokobori E, Uchiyama M, Matsuda K. 2012. Gonadotropin-releasing hormone 2 suppresses food intake in the zebrafish, Danio rerio. Frontiers in Endocrinology 3. DOI: 10.3389/fendo.2012.00122

Nudo RJ, Masterton RB. 1989. Descending pathways to the spinal cord: II. Quantitative study of the tectospinal tract in 23 mammals. Journal of Comparative Neurology 286:96–119. DOI: 10.1002/cne.902860107

Nudo RJ, Masterton RB. 1988a. Descending pathways to the spinal cord: A comparative study of 22 mammals. Journal of Comparative Neurology 277:53–79. DOI: 10.1002/cne.902770105

Nudo RJ, Masterton RB. 1988b. Descending pathways to the spinal cord: A comparative study of 22 mammals. Journal of Comparative Neurology 277:53–79. DOI: 10.1002/cne.902770105

Nyberg-Hansen R. 1965. SITES AND MODE OF TERMINATION OF RETICULO-SPINAL FIBERS IN THE CAT. AN EXPERIMENTAL STUDY WITH SILVER IMPREGNATION METHODS. The Journal of Comparative Neurology 124:71–99. DOI: 10.1002/cne.901240107, PMID: 14304275

Ocaña FM, Suryanarayana SM, Saitoh K, Kardamakis AA, Capantini L, Robertson B, Grillner S. 2015. The Lamprey Pallium Provides a Blueprint of the Mammalian Motor Projections from Cortex. Current Biology 25:413–423. DOI: 10.1016/j.cub.2014.12.013

Oka Y, Satou M, Ueda K. 1986. Descending pathways to the spinal cord in the himé salmon (landlocked red salmon, Oncorhynchus nerka). The Journal of Comparative Neurology 254:91–103. DOI: 10.1002/cne.902540108, PMID: 3805356

Olson I, Suryanarayana SM, Robertson B, Grillner S. 2017. Griseum centrale, a homologue of the periaqueductal gray in the lamprey. IBRO Reports 2:24–30. DOI: 10.1016/j.ibror.2017.01.001

Opdam P, Kemali M, Nieuwenhuys R. 1976. Topological analysis of the brain stem of the frogs Rana esculenta and Rana catesbeiana. Journal of Comparative Neurology 165:307–331. DOI: 10.1002/cne.901650304

Orger MB, Kampff AR, Severi KE, Bollmann JH, Engert F. 2008. Control of visually guided behavior by distinct populations of spinal projection neurons. Nature Neuroscience 11:327–333. DOI: 10.1038/nn2048

Oti T, Satoh K, Uta D, Nagafuchi J, Tateishi S, Ueda R, Takanami K, Young LJ, Galione A, Morris JF, Sakamoto T, Sakamoto H. 2021. Oxytocin Influences Male Sexual Activity via Non-synaptic Axonal Release in the Spinal Cord. Current Biology 31:103–114.e5. DOI: 10.1016/j.cub.2020.09.089

Pansera L, Mhalhel K, Cavallaro M, Aragona M, Laurà R, Levanti M, Guerrera MC, Abbate F, Germanà A, Montalbano G. 2025. Zebrafish as an Integrative Model for Central Nervous System Research: Current Advances and Translational Perspectives. Life 15. DOI: 10.3390/life15111751

Papez JW. 1926. Reticulo-spinal tracts in the cat. Marchi method. Journal of Comparative Neurology 41:365–399. DOI: 10.1002/cne.900410113

Pompeiano O. 1973. Reticular Formation. In: Albe-Fessard D, Andres KH, Bates JAV, Besson JM, Brown AG, Burgess PR, Darian-Smith I, v. Düring M, Gordon G, Hensel H, Jones E, Libet B, Oscarsson O, Perl ER, Pompeiano O., Powell TPS, Réthelyi M, Schmidt RF, Semmes J, Skoglund S, Szentágothai J, Towe AL, Wall PD, Werner G, Whitsel BL, Zotterman Y, Iggo A (Eds). Somatosensory System, Handbook of Sensory Physiology. Springer. p. 381–488. DOI: 10.1007/978-3-642-65438-1_13

Posit team. 2025. RStudio: Integrated Development Environment for R.

Prasada Rao PD, Jadhao AG, Sharma SC. 1987a. Descending projection neurons to the spinal cord of the goldfish, Carassius auratus. The Journal of Comparative Neurology 265:96–108. DOI: 10.1002/cne.902650107, PMID: 2826554

Prasada Rao PD, Jadhao AG, Sharma SC. 1987b. Descending projection neurons to the spinal cord of the goldfish, Carassius auratus. The Journal of Comparative Neurology 265:96–108. DOI: 10.1002/cne.902650107, PMID: 2826554

Preibisch S, Saalfeld S, Tomancak P. 2009. Globally optimal stitching of tiled 3D microscopic image acquisitions. Bioinformatics 25:1463–1465. DOI: 10.1093/bioinformatics/btp184

Priest MF, Freda SN, Rieth IJ, Badong D, Dumrongprechachan V, Kozorovitskiy Y. 2023. Peptidergic and functional delineation of the Edinger-Westphal nucleus. Cell reports 42:112992. DOI: 10.1016/j.celrep.2023.112992, PMID: 37594894

Pujala A, Koyama M. 2019. Chronology-based architecture of descending circuits that underlie the development of locomotor repertoire after birth. eLife 8:e42135. DOI: 10.7554/eLife.42135

Randlett O, Wee CL, Naumann EA, Nnaemeka O, Schoppik D, Fitzgerald JE, Portugues R, Lacoste AMB, Riegler C, Engert F, Schier AF. 2015. Whole-brain activity mapping onto a zebrafish brain atlas. Nature Methods 12:1039–1046. DOI: 10.1038/nmeth.3581

Ronan M. 1989. Origins of the descending spinal projections in petromyzontid and myxinoid agnathans. The Journal of Comparative Neurology 281:54–68. DOI: 10.1002/cne.902810106, PMID: 2925902

Ronan MC, Northcutt RG. 1985. The origins of descending spinal projections in lepidosirenid lungfishes. Journal of Comparative Neurology 241:435–444. DOI: 10.1002/cne.902410404

Rubinson K. 2008. Projections of the Tectum Opticum of the Frog; pp. 529–544. Brain Behavior and Evolution 1:529–544. DOI: 10.1159/000125524

Ruta V, Datta SR, Vasconcelos ML, Freeland J, Looger LL, Axel R. 2010. A dimorphic pheromone circuit in Drosophila from sensory input to descending output. Nature 468:686–690. DOI: 10.1038/nature09554, PMID: 21124455

Ryczko D, Cone JJ, Alpert MH, Goetz L, Auclair F, Dubé C, Parent M, Roitman MF, Alford S, Dubuc R. 2016. A descending dopamine pathway conserved from basal vertebrates to mammals. Proceedings of the National Academy of Sciences 113:E2440–E2449. DOI: 10.1073/pnas.1600684113

Sánchez-Camacho C, Marín O, Ten Donkelaar HJ, González A. 2001. Descending supraspinal pathways in amphibians. I. A dextran amine tracing study of their cells of origin. Journal of Comparative Neurology 434:186–208. DOI: 10.1002/cne.1172

Severi KE, Portugues R, Marques JC, O’Malley DM, Orger MB, Engert F. 2014. Neural control and modulation of swimming speed in the larval zebrafish. Neuron 83:692–707. DOI: 10.1016/j.neuron.2014.06.032, PMID: 25066084

Sharples SA, Koblinger K, Humphreys JM, Whelan PJ. 2014. Dopamine: a parallel pathway for the modulation of spinal locomotor networks. Frontiers in Neural Circuits 8:55. DOI: 10.3389/fncir.2014.00055, PMID: 24982614

Sinnamon HM, Benaur M. 1997. GABA injected into the anterior dorsal tegmentum (ADT) of the midbrain blocks stepping initiated by stimulation of the hypothalamus. Brain Research 766:271–275. DOI: 10.1016/s0006-8993(97)00734-8, PMID: 9359615

Skinner RD, Garcia-Rill E, Griffin S, Nelson R, Fitzgerald JA. 1984. Interstitial nucleus of Cajal (INC) projections to the region of Probst’s tract. Brain Research Bulletin 13:613–621. DOI: 10.1016/0361-9230(84)90192-8

Smeets WJ, Timerick SJ. 1981a. Cells of origin of pathways descending to the spinal cord in two chondrichthyans, the shark Scyliorhinus canicula and the ray Raja clavata. The Journal of Comparative Neurology 202:473–491. DOI: 10.1002/cne.902020403, PMID: 7298910

Smeets WJ, Timerick SJ. 1981b. Cells of origin of pathways descending to the spinal cord in two chondrichthyans, the shark Scyliorhinus canicula and the ray Raja clavata. The Journal of Comparative Neurology 202:473–491. DOI: 10.1002/cne.902020403, PMID: 7298910

Smith CM, Shen P-J, Banerjee A, Bonaventure P, Ma S, Bathgate RAD, Sutton SW, Gundlach AL. 2010. Distribution of relaxin-3 and RXFP3 within arousal, stress, affective, and cognitive circuits of mouse brain. Journal of Comparative Neurology 518:4016–4045. DOI: 10.1002/cne.22442

Sofroniew N, Lambert T, Bokota G, Nunez-Iglesias J, Sobolewski P, Sweet A, Gaifas L, Evans K, Burt A, Doncila Pop D, Yamauchi K, Weber Mendonça M, Rodríguez-Guerra J, Liu L, Buckley G, Vierdag W-M, Anderson A, Monko T, Willing C, Royer L, Can Solak A, Harrington KIS, Abramo J, Ahlers J, Althviz Moré D, Amsalem O, Andò E, Annex A, Aronssohn C, Balzaretti F, Boone P, Bragantini J, Bunten D, Bussonnier M, Caporal C, Coccimiglio I, Čočková Z, Eglinger J, Eisenbarth A, Freeman J, Fukai T. Y, Gohlke C, Gunalan K, Halchenko YO, Har-Gil H, Harfouche M, Hilsenstein V, Hutchings K, Kozar R, Lauer J, Lichtner G, Liu H, Liu Z, Lowe A, Marconato L, Martin S, McGovern A, Migas L, Miller N, Miñano S, Muñoz H, Müller J-H, Nauroth-Kreß C, Obenhaus HA, Palecek D, Pape C, Perlman E, Theart RP, Pevey K, Peña-Castellanos G, Pierré A, Pinto D, Rodríguez-Reza CM, Ross D, Russell CT, Ryan J, Selzer G, Smith M, Smith P, Sofiiuk K, Soltwedel J, Stansby D, Vanaret J, Wadhwa P, Weigert M, Windhager J, Winston P, Yu Q, Zhao R, Witz G. 2026. napari: a multi-dimensional image viewer for Python. DOI: 10.5281/zenodo.18374344

Spikol ED, Cheng J, Macurak M, Subedi A, Halpern ME. 2024. Genetically defined nucleus incertus neurons differ in connectivity and function. eLife 12:RP89516. DOI: 10.7554/eLife.89516

Swain GP, Snedeker JA, Ayers J, Selzer ME. 1993. Cytoarchitecture of spinal-projecting neurons in the brain of the larval sea lamprey. The Journal of Comparative Neurology 336:194–210. DOI: 10.1002/cne.903360204, PMID: 8245215

Swanson LW, Sawchenko PE. 1980. Paraventricular Nucleus:A Site for the Integration of Neuroendocrine and Autonomic Mechanisms. Neuroendocrinology 31:410–417. DOI: 10.1159/000123111

Tay TL, Ronneberger O, Ryu S, Nitschke R, Driever W. 2011. Comprehensive catecholaminergic projectome analysis reveals single-neuron integration of zebrafish ascending and descending dopaminergic systems. Nature Communications 2:171. DOI: 10.1038/ncomms1171

Teclemariam-Mesbah R, Kalsbeek A, Buijs RM, Pévet P. 1997. Oxytocin innervation of spinal preganglionic neurons projecting to the superior cervical ganglion in the rat. Cell and Tissue Research 287:481–486. DOI: 10.1007/s004410050772, PMID: 9023079

Ten Donkelaar HJ. 2020. Clinical Neuroanatomy: Brain Circuitry and Its Disorders. Springer International Publishing. DOI: 10.1007/978-3-030-41878-6

ten Donkelaar HJ. 2009. Evolution of Motor Systems: Corticospinal, Reticulospinal, Rubrospinal and Vestibulospinal Systems. In: Binder MD, Hirokawa N, Windhorst U (Eds). Encyclopedia of Neuroscience. Springer. p. 1248–1254. DOI: 10.1007/978-3-540-29678-2_3131

ten Donkelaar HJ. 1988. Evolution of the red nucleus and rubrospinal tract. Behavioural Brain Research 28:9–20. DOI: 10.1016/0166-4328(88)90072-1, PMID: 3289562

Ten Donkelaar HJ. 1982a. Organization of descending pathways to the spinal cord in amphibians and reptiles. Progress in Brain Research 57:25–67. DOI: 10.1016/s0079-6123(08)64123-0, PMID: 7156397

Ten Donkelaar HJ. 1982b. Organization of descending pathways to the spinal cord in amphibians and reptiles. Progress in Brain Research 57:25–67. DOI: 10.1016/s0079-6123(08)64123-0, PMID: 7156397

Ten Donkelaar HJ. 1976. Descending pathways from the brain stem to the spinal cord in some reptiles. I. Origin. The Journal of Comparative Neurology 167:421–442. DOI: 10.1002/cne.901670403, PMID: 1270628

ten Donkelaar HJ, de Boer-van Huizen R, Schouten FTM, Eggen SJH. 1981a. Cells of origin of descending pathways to the spinal cord in the clawed toad (Xenopus laevis). Neuroscience 6:2297–2312. DOI: 10.1016/0306-4522(81)90019-1

ten Donkelaar HJ, de Boer-van Huizen R, Schouten FTM, Eggen SJH. 1981b. Cells of origin of descending pathways to the spinal cord in the clawed toad (Xenopus laevis). Neuroscience 6:2297–2312. DOI: 10.1016/0306-4522(81)90019-1

ten Donkelaar HJ, Kusuma A, de Boer-Van Huizen R. 1980a. Cells of origin of pathways descending to the spinal cord in some quadrupedal reptiles. The Journal of Comparative Neurology 192:827–851. DOI: 10.1002/cne.901920413, PMID: 7419757

ten Donkelaar HJ, Kusuma A, de Boer-Van Huizen R. 1980b. Cells of origin of pathways descending to the spinal cord in some quadrupedal reptiles. The Journal of Comparative Neurology 192:827–851. DOI: 10.1002/cne.901920413, PMID: 7419757

Thiele TR, Donovan JC, Baier H. 2014. Descending control of swim posture by a midbrain nucleus in zebrafish. Neuron 83:679–691. DOI: 10.1016/j.neuron.2014.04.018, PMID: 25066082

Tohyama M, Sakai K, Salvert D, Touret M, Jouvet M. 1979. Spinal projections from the lower brain stem in the cat as demonstrated by the horseradish peroxidase technique. I. Origins of the reticulospinal tracts and their funicular trajectories. Brain Research 173:383–403. DOI: 10.1016/0006-8993(79)90237-3

Topilko T, Diaz SL, Pacheco CM, Verny F, Rousseau CV, Kirst C, Deleuze C, Gaspar P, Renier N. 2022. Edinger-Westphal peptidergic neurons enable maternal preparatory nesting. Neuron 110:1385–1399.e8. DOI: 10.1016/j.neuron.2022.01.012, PMID: 35123655

Torvik A, Brodal A. 1957. The origin of reticulospinal fibers in the cat; an experimental study. The Anatomical Record 128:113–137. DOI: 10.1002/ar.1091280110, PMID: 13458831

Urasaki A, Asakawa K, Kawakami K. 2008. Efficient transposition of the Tol2 transposable element from a single-copy donor in zebrafish. Proceedings of the National Academy of Sciences 105:19827–19832. DOI: 10.1073/pnas.0810380105

VanderHorst VGJM, Ulfhake B. 2006. The organization of the brainstem and spinal cord of the mouse: Relationships between monoaminergic, cholinergic, and spinal projection systems. Journal of Chemical Neuroanatomy 31:2–36. DOI: 10.1016/j.jchemneu.2005.08.003

Verstegen AMJ, Vanderhorst V, Gray PA, Zeidel ML, Geerling JC. 2017. Barrington’s nucleus: Neuroanatomic landscape of the mouse “pontine micturition center.” Journal of Comparative Neurology 525:2287–2309. DOI: 10.1002/cne.24215

Walter BL, Shaikh AG. 2014. Midbrain. Encyclopedia of the Neurological Sciences. Elsevier. p. 28–33. DOI: 10.1016/B978-0-12-385157-4.01161-1

Wang X, Liu Y, Li X, Zhang Z, Yang H, Zhang Yu, Williams PR, Alwahab NSA, Kapur K, Yu B, Zhang Yiming, Chen M, Ding H, Gerfen CR, Wang KH, He Z. 2017. Deconstruction of Corticospinal Circuits for Goal-Directed Motor Skills. Cell 171:440–455.e14. DOI: 10.1016/j.cell.2017.08.014, PMID: 28942925

Wang Z, Romanski A, Mehra V, Wang Y, Brannigan M, Campbell BC, Petsko GA, Tsoulfas P, Blackmore MG. 2022. Brain-wide analysis of the supraspinal connectome reveals anatomical correlates to functional recovery after spinal injury. eLife 11:e76254. DOI: 10.7554/eLife.76254

Warner FJ. 1947. The Diencephalon and Midbrain of the American Rattlesnake (Crotalus adamanteus). Proceedings of the Zoological Society of London 116:531–550. DOI: 10.1111/j.1096-3642.1947.tb00133.x

Watson C, Bartholomaeus C, Puelles L. 2019a. Time for Radical Changes in Brain Stem Nomenclature—Applying the Lessons From Developmental Gene Patterns. Frontiers in Neuroanatomy 13.

Watson C, Bartholomaeus C, Puelles L. 2019b. Time for Radical Changes in Brain Stem Nomenclature-Applying the Lessons From Developmental Gene Patterns. Frontiers in Neuroanatomy 13:10. DOI: 10.3389/fnana.2019.00010, PMID: 30809133

Wee CL, Nikitchenko M, Wang W-C, Luks-Morgan SJ, Song E, Gagnon JA, Randlett O, Bianco IH, Lacoste AMB, Glushenkova E, Barrios JP, Schier AF, Kunes S, Engert F, Douglass AD. 2019. Zebrafish oxytocin neurons drive nocifensive behavior via brainstem premotor targets. Nature Neuroscience 22:1477–1492. DOI: 10.1038/s41593-019-0452-x

White RM, Sessa A, Burke C, Bowman T, LeBlanc J, Ceol C, Bourque C, Dovey M, Goessling W, Burns CE, Zon LI. 2008a. Transparent adult zebrafish as a tool for in vivo transplantation analysis. Cell Stem Cell 2:183–189. DOI: 10.1016/j.stem.2007.11.002, PMID: 18371439

White RM, Sessa A, Burke C, Bowman T, LeBlanc J, Ceol C, Bourque C, Dovey M, Goessling W, Burns CE, Zon LI. 2008b. Transparent adult zebrafish as a tool for in vivo transplantation analysis. Cell stem cell 2:183–189. DOI: 10.1016/j.stem.2007.11.002, PMID: 18371439

Winter CC, Wang KH, He Z. 2025. From thought to action: The organization of spinal projecting neurons. Cell Reports 44:116153. DOI: 10.1016/j.celrep.2025.116153

Wolters JG, -Van Huizen RDB, Ten Donkelaar HJ. 1982. Funicular Trajectories of Descending Brain Stem Pathways in a Lizard(Varanus exanthematicus). In: Kuypers HGJM, Martin GF (Eds). Progress in Brain Research. Elsevier. p. 69–78. DOI: 10.1016/S0079-6123(08)64124-2

Wullimann MF, Mokayes N, Shainer I, Kuehn E, Baier H. 2023. Genoarchitectonics of the larval zebrafish diencephalon. The Journal of comparative neurology. DOI: 10.1002/cne.25549, PMID: 37983970

Xia W, Smith O, Zmora N, Xu S, Zohar Y. 2014. Comprehensive analysis of GnRH2 neuronal projections in zebrafish. Scientific Reports 4:3676. DOI: 10.1038/srep03676, PMID: 24419253

Yamamoto N, Nakayama T, Hagio H. 2017. Descending pathways to the spinal cord in teleosts in comparison with mammals, with special attention to rubrospinal pathways. Development, Growth & Differentiation 59:188–193. DOI: 10.1111/dgd.12355

Yamamoto N, Oka Y, Amano M, Aida K, Hasegawa Y, Kawashima S. 1995a. Multiple gonadotropin-releasing hormone (GnRH)-immunoreactive systems in the brain of the dwarf gourami, Colisa lalia: immunohistochemistry and radioimmunoassay. The Journal of Comparative Neurology 355:354–368. DOI: 10.1002/cne.903550303, PMID: 7636018

Yamamoto N, Oka Y, Amano M, Aida K, Hasegawa Y, Kawashima S. 1995b. Multiple gonadotropin-releasing hormone (GnRH)-immunoreactive systems in the brain of the dwarf gourami, Colisa lalia: immunohistochemistry and radioimmunoassay. The Journal of Comparative Neurology 355:354–368. DOI: 10.1002/cne.903550303, PMID: 7636018

Yamamoto N, Oka Y, Yoshimoto M, Sawai N, Albert JS, Ito H. 1998. Gonadotropin-releasing hormone neurons in the gourami midbrain: a double labeling study by immunocytochemistry and tracer injection. Neuroscience Letters 240:50–52. DOI: 10.1016/s0304-3940(97)00906-3, PMID: 9488172

Yáñez J, Eguiguren MH, Anadón R. 2024. Neural connections of the torus semicircularis in the adult Zebrafish. Journal of Comparative Neurology 532:e25586. DOI: 10.1002/cne.25586

Zhao H, Liu J, Shao Y, Feng X, Zhao B, Sun L, Liu Y, Zeng L, Li X, Yang H, Duan S, Yu Y. 2024. Control of defensive behavior by the nucleus of Darkschewitsch GABAergic neurons. National Science Review 11:nwae082. DOI: 10.1093/nsr/nwae082, PMID: 38686177

